# The full spectrum of OCT1 (SLC22A1) mutations bridges transporter biophysics to drug pharmacogenomics

**DOI:** 10.1101/2023.06.06.543963

**Authors:** Sook Wah Yee, Christian Macdonald, Darko Mitrovic, Xujia Zhou, Megan L. Koleske, Jia Yang, Dina Buitrago Silva, Patrick Rockefeller Grimes, Donovan Trinidad, Swati S. More, Linda Kachuri, John S. Witte, Lucie Delemotte, Kathleen M. Giacomini, Willow Coyote-Maestas

## Abstract

Membrane transporters play a fundamental role in the tissue distribution of endogenous compounds and xenobiotics and are major determinants of efficacy and side effects profiles. Polymorphisms within these drug transporters result in inter-individual variation in drug response, with some patients not responding to the recommended dosage of drug whereas others experience catastrophic side effects. For example, variants within the major hepatic Human organic cation transporter OCT1 (SLC22A1) can change endogenous organic cations and many prescription drug levels. To understand how variants mechanistically impact drug uptake, we systematically study how all known and possible single missense and single amino acid deletion variants impact expression and substrate uptake of OCT1. We find that human variants primarily disrupt function via folding rather than substrate uptake. Our study revealed that the major determinants of folding reside in the first 300 amino acids, including the first 6 transmembrane domains and the extracellular domain (ECD) with a stabilizing and highly conserved stabilizing helical motif making key interactions between the ECD and transmembrane domains. Using the functional data combined with computational approaches, we determine and validate a structure-function model of OCT1s conformational ensemble without experimental structures. Using this model and molecular dynamic simulations of key mutants, we determine biophysical mechanisms for how specific human variants alter transport phenotypes. We identify differences in frequencies of reduced function alleles across populations with East Asians vs European populations having the lowest and highest frequency of reduced function variants, respectively. Mining human population databases reveals that reduced function alleles of OCT1 identified in this study associate significantly with high LDL cholesterol levels. Our general approach broadly applied could transform the landscape of precision medicine by producing a mechanistic basis for understanding the effects of human mutations on disease and drug response.

## INTRODUCTION

For most drugs the relationship between dose and response varies widely between individuals, due to differential absorption, metabolism, and transport. Membrane-spanning transporters play a critical role in pharmacokinetics of many drugs, especially more polar molecules, by allowing them to pass the hydrophobic lipid bilayer. The human genome encodes a vast array of drug transporters, including ATP Binding Cassette (ABC) and solute carriers (SLCs) (Ferrada and Superti-Furga, 2022). Among these, the organic cation transporter subgroup of the SLC22 family (OCTs) is a key group of polyspecific transporters that are expressed in the liver, kidney, and intestine. With their tissue-specific expression and broad substrate selectivity, OCTs play major roles in the uptake of endogenous molecules and xenobiotics (Gorboulev *et al*., 1997; Zhang *et al*., 1997; Yee and Giacomini, 2021). OCT1 transports a range of cations (e.g., MPP^+^, TEA), endogenous molecules (e.g., thiamine (Chen *et al*., 2014) and serotonin (Haberkorn, Fromm and König, 2021)), and prescription drugs (e.g., metformin (Shu *et al*., 2007) and morphine (Meyer *et al*., 2019)).

Reduced OCT1 function results in increased plasma drug levels, altered therapeutic windows (Matthaei *et al*., 2016; Stamer *et al*., 2016; Tzvetkov *et al*., 2018), and changes to endogenous metabolite and lipid levels (Long *et al*., 2017; Hoffmann *et al*., 2018; Liang *et al*., 2018; Yousri *et al*., 2018). While there is ample evidence of OCT1 polymorphisms playing roles in modulating drug responses, thus far only 39 of the 100s of known common and rare variants have been characterized in any way. Furthermore, there are well over 10,000 possible variants within OCT1 which could and likely are contributing to differences in therapeutic and adverse drug response among people. More broadly, similar issues plague our understanding of other SLC and ABC drug transporters with substantial genetic variation, but little connection to function. To understand how humans display such diverse responses to therapeutics we need comprehensive models of how genetic variation impacts drug transporters. Such models must also be mechanistic because some variants impact the uptake of all drugs such as when protein folding is disrupted whereas other variants may affect substrate specificity. Therefore, an understanding for how a variant impacts drug uptake requires models that can predict how mutations affect the biophysics of folding, the structural biology of uptake, and the genomics of human physiology.

In this exploratory investigation, we take the first step towards this goal by comprehensively studying how all single mutations and amino acid deletions alter OCT1 folding, function, and human physiology. In particular, we determined the impact on expression and function of 11,213 variants of OCT1. Based on the expression data, we identify which mutations alter folding and which regions drive folding in the SLC22 fold family. Using our functional data combined with predictions of conformational ensembles of OCT1 with and without ligands we can build a structure-function model for OCT1 without experimental structures. Using this model as a starting point we identified how mutations biophysically disrupt OCT1’s function. Finally, by comparing mutational data with large scale population and biomedical databases, we determine how poor function variants are distributed across different global populations, impact human physiology, and drug response

## Results

### An expression and function deep mutational scan of OCT1

In this study our goal was to develop a mechanistic understanding for how polymorphisms alter OCT1 function. To this end, we conducted a deep mutational scanning (DMS) experiment in which we experimentally determine the effect of a comprehensive library of single missense and deletion variants in pooled genetic screens (Fowler and Fields, 2014). Mutations can impact whether OCT1 can express and fold, whether they eliminate function completely or whether they have substrate specific effects. To identify the mechanistic basis by which mutations alter a gene, multi-parametric assays can be used that decompose ‘fitness’ into more granular protein properties such as expression and protein activity (Amorosi *et al*., 2021; Markin *et al*., 2021; Coyote-Maestas *et al*., 2022; Faure *et al*., 2022). For OCT1 we developed two approaches based on the uptake of a cytotoxic substrate and the fluorescence of OCT1 to understand whether mutations alter function and folding. To monitor the impact of mutations on function within a negative selection screen, we used a cytotoxic substrate, SM73, which is a cisplatin analog that we had previously developed to kill cancer cells in an OCT1-dependent manner with sensitivity to variants effects (Figure 1A-B). To measure how mutations impact expression, we C-terminally fluorescently tagged OCT1 with a split fluorescent protein, mNeonGreen, component and could differentiate variant impacts based on fluorescence in FACS-seq experiments. sort variants (Figure 1C-D) (Matreyek et al, 2018). Using these assays on a mutational library in HEK293T cells, we measured how nearly all single missense and single codon deletion variants impacts OCT1 function and expression (Matreyek *et al*., 2020; Macdonald *et al*., 2023) (Figures 1D-E, 2, Supp Fig 1-3). We find that the assays are highly reproducible and generate high quality data (R^2^ 0.781-0.878, cytotoxicity; R^2^ 0.852-0.871, VAMP-seq; Fig 2, Supp Figs 4-7).

**Figure 1.**
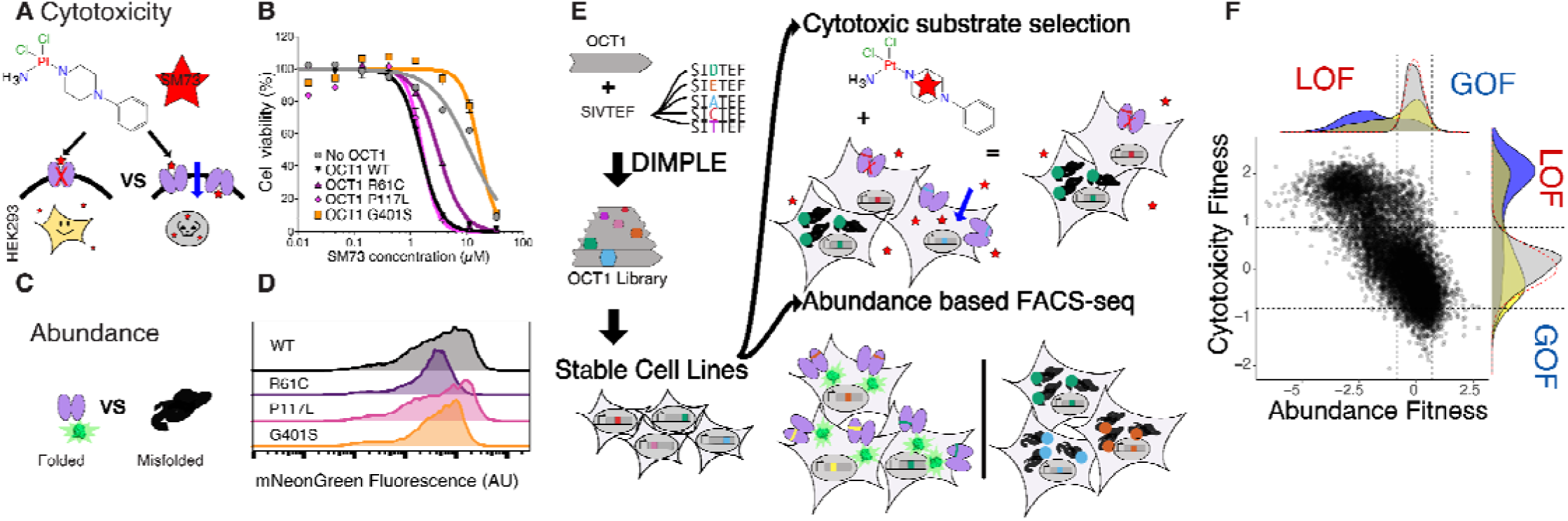
Workflow for multiparametric deep mutational scan of OCT1. **(A)** Cytotoxic OCT1 substrates such as SM73 enable negative selection screens for OCT1 function in HEK293 cells. **(B)** OCT1 variants have different sensitivity to Cisplatin analog SM85, allowing a fitness gradient. (**C)** A split fluorescent protein-based readout for variant abundance can be used to distinguish folded vs misfolded forms of OCT1 as demonstrated (**D**) in flow cytometry experiments where loss of transport variants from cytotoxicity screening have diminished fluorescence. **(E)** We generated an OCT1 deep mutational scanning library with our DIMPLE protocol, produced stable cell lines, and conducted parallel abundance and cytotoxicity selection screens to determine the functional impacts of variants, yielding **(F)** a multiparametric fitness landscape for 11,213 OCT1 mutants: x-axis, abundance score, and y-axis, cytotoxicity scores, with density plots indicating classes of mutations including synonymous (grey), missense (yellow), and single codon deletion (purple). Cutoffs for loss and gain of function for both phenotypes are indicated by a dotted line, which were determined using a 2 standard deviations from normal distribution fit upon synonymous distributions.

**Figure 2.**
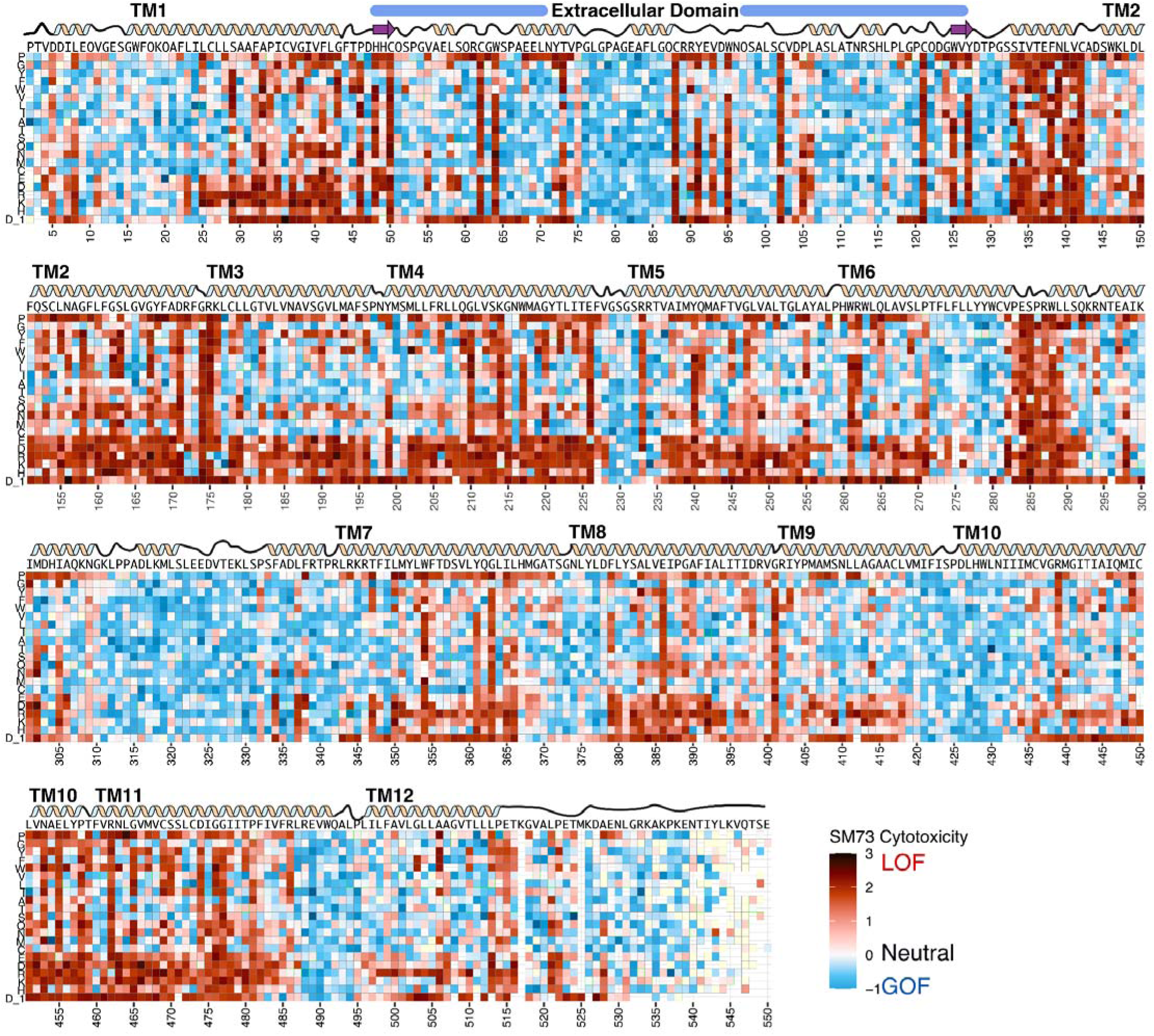
Heatmap of OCT1 cytotoxicity deep mutational scan. Cytotoxicity screen fitness effects depicted as a heatmap, with (x-axis) residue position versus (y-axis) mutation identity. In this assay, *loss* of transport activity (relative to wildtype) reduces the uptake of a cytotoxic substrate, thus increasing abundance (positive score, more red) while *gain* of transport *increases* substrate uptake, which *decreases* abundance (negative score, more blue). Above, wildtype sequence and representation of secondary structure elements of OCT1. Missing data in light yellow.

The cytotoxicity assay is a readout for whether OCT1 is functional which is a combination of expression and substrate uptake. To determine whether expression or substrate uptake is the major determinant of function, we compared scores and found that cytotoxicity scores correlate strongly with abundance implying mutations primarily impact expression. That expression primarily determines function follows previous studies in more limited mutational screens in transporters as well as DMS experiments in channels and GPCRs, which find folding is the most common effect of mutations (Penn *et al*., 2020; Young *et al*., 2021; Coyote-Maestas *et al*., 2022; Koleske *et al*., 2022; McKee *et al*., 2022; Muhammad *et al*., 2023). To test which screen better represents overall function, we compared conservation to these scores and unsurprisingly we find conservation has a stronger correlation to the cytotoxicity than to the abundance data (0.59 vs 0.48 Spearman correlation coefficient). Cytotoxicity would likely be closer to the presumed phenotype under selection within evolution, transport function, than abundance even if abundance explains most of the variance. This OCT1 DMS experiment allows us to determine that variants primarily impact expression while the cytotoxicity assay most closely resembles evolution.

### Transmembrane helices 1-6 and the extracellular domain drive stability and folding

From comparing abundance and cytotoxicity scores we discovered that mutations primarily impact OCT1 function through changes in protein expression. Because the primary determinant of expression are folding, we sought to determine how mutations mechanistically alter folding. To identify which residues underlie folding and stability, we calculated a folding importance score defined as the summed absolute value of VAMP-seq scores at each position across all loss of function mutations (see Methods). Interestingly, critical residues for folding are enriched within the first 300 amino acids of OCT1 and not the C-terminal amino acid residues (Figure 3A, Supp). Folding of proteins tends to be driven by stabilizing interactions between amino acids including the packing of hydrophobic residues, disulfide bonds, and ionic interactions. To test that the specific physicochemistry of these residues are important within folding critical regions, we compared how changing between amino acid types impacted residues with high vs low stability importance. We find cysteines, aromatics, and negative residues with high stability importance are particularly sensitive to mutations implying that cysteines are forming disulfides, aromatics are packing in hydrophobic cores, and charged residues forming ionic bonds. Strikingly, these same residue types with low abundance scores are dramatically less sensitive to physicochemical changes (Figure 3H-J).

**Figure 3.**
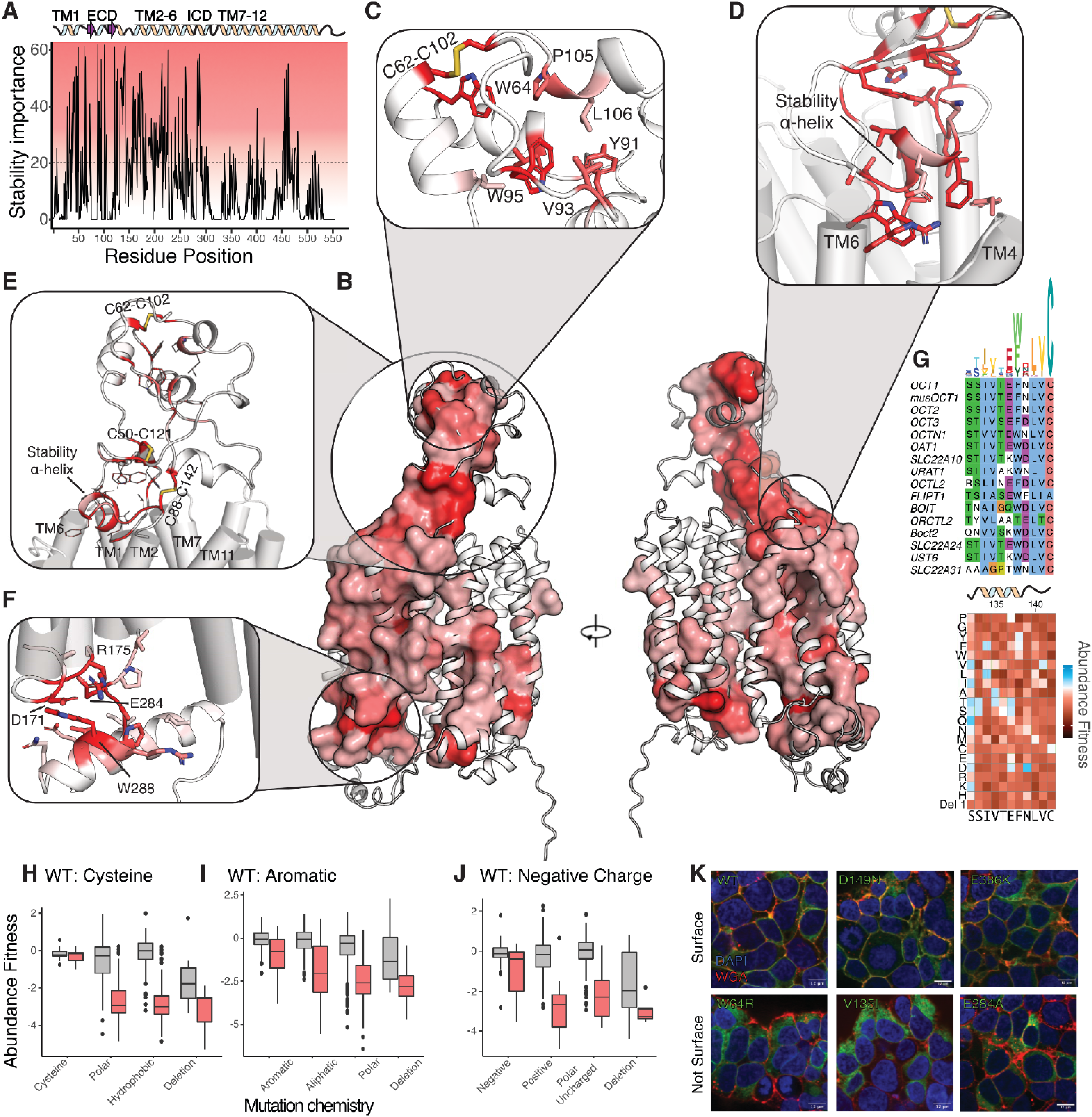
OCT1 folding determinants revealed by mutational scan. **(A)** Stability importance scores plotted across OCT1 positions (with a simplified architecture above) suggests that TM1-6 are the primary determinants of stability and folding. Dotted line indicates threshold for identifying stability determinants. **(B)** Mapping the stability determinant scores mapped onto the OCT1 AlphaFold2 model reveals a dense network of residues connecting the cytosolic transmembrane bundles with the extracellular domain. Residues with stability importance scores above the cutoff in **A** are shown as a surface. (**C)** In the extracellular domain, the distinct fold has a hydrophobic core composed of aromatics and disulfide bonds which have strong impacts on stability. (**D**-**E)** A dense network of disulfides, aromatics, and other residues connect the transmembrane domains to the ECD via a stabilizing alpha helix, which interacts with the tops of TMs 4 and 6. **(F)** A series of charged residues are highly important for stability and folding among the cytoplasmic termini of TM1-6. (**G**) Multiple sequence alignment of the stability helix with a sequence logo above with height corresponding to information content. Below: experimental VAMP-seq scores for the region of OCT1. **(H-J)** Positions with high folding importance (red boxplots) show more sensitivity to changes of physical chemistry than other positions (gray boxplot): the abundance scores of WT cysteine, aromatic, or negatively charged residues show larger effects of changes to physical chemistry in regions of higher importance, characteristic of folding determinants. **(K)** Confocal microscopy validation of trafficking phenotypes of variants from abundance screen. Using EGFP-tagged OCT1 (green), cellular localization and expression are determined using nuclear stain (DAPI, blue) and cell surface stain (WGA-Alexa Fluor 647, red). Variants with low abundance identified in our screen (W64R, V135I, E284A) have commensurate low surface trafficking whereas WT and WT-like variants (D149N and E386K) are properly expressed and trafficked to the surface.

We were especially interested in investigating why residues important to folding are enriched in the first 300 amino acids as well as learning how amino acids are interacting to drive folding. To do so, we mapped these onto the OCT1 protein structure predicted by AlphaFold2 because there are no experimental structures of OCT1 available with a fully resolved extracellular domain. The AlphaFold2 prediction has high confidence within structured regions that are highly similar to the resolved portions of solved experimental structures of OCT1 and OCT3 (Supp Fig 8, Supp Table 1). As with other Major Facilitator Superfamily (MFS) SLC transporters, OCT1 is made up of two six-transmembrane domain bundles, which are typically termed the N– and C-bundles. Distinct to OCT1 and other SLC22’s is an extracellular domain (ECD) between TMD1 and TMD2, which has not been experimentally resolved but exists within SLC22 subfamily AlphaFold2 models. In fact, it appears that this large ECD exists only within the SLC22 family with no similar folds detected across experimental or predicted structures using the structure-based protein homology search server Dali (Holm, 2022). Using the structure as a scaffold to map the folding importance scores reveals that primarily folding is driven by the N-terminal 6 transmembrane bundle, the ECD, and a small cluster of residues that interact at the bottom of the second 6 transmembrane domain bundle (Fig 3A-B, Supp Fig 9).

The ECD starts immediately after the first transmembrane domain and therefore the biogenesis of OCT1 will be dependent on proper folding of the ECD. The ECD is composed of a mixture of alpha helices and beta sheets with large loops (Figures 3B-E). The distal pair of helices interact with a loop, forming a hydrophobic core with many folding critical residues, including aromatic (W64, Y91), hydrophobic, (V93, L106), and cysteines forming a disulfide bond (C62-C102) (Figure 3C). All the mutations within this motif are extremely sensitive to changes in physichochemistry. While OCT1 folding is dependent on this motif, it does not appear to be strongly conserved across SLC22 transporters. For example, URAT1 does not contain the disulfide bond nor enrichment of aromatics, implying that conservation-based machine models of loss of function such as EVE would not pick up mutations in this region (Frazer *et al*., 2021). Previous studies had identified putative glycosylation sites at ECD residues (71, 72, 96, 98, and 99) based on physicochemistry and conservation, however in our screens, we find these residues are quite mutable to non-glycosylated residues implying they are not necessary for expression and function (Burckhardt and Wolff, 2000).

An adjacent network of residues connects the distal region to the transmembrane domain of OCT1, comprising a pair of double-stranded antiparallel beta-sheets among a pair of disulfide bonds, which do not allow mutations besides (C50-C121 and C88-C142) abutting a pair α-helices interacting with alternating TMH1-6 or TMH 7-12 (Figure 3D-E). Each of these secondary structures are extremely sensitive to mutations and the disulfide bonds do not allow non-cysteine substitutions. We call the TMH1-6 interacting alpha helix and following residues (133-SIVTEFNLVC) the *stability* α*-helix* makes key interactions between TMs 1, 3, and 6 and the ECD, is enriched in stability-determining positions, and is extremely sensitive to substitutions that change amino acid physicochemistry. The stability helix and several residues afterward and contain a charged residue that likely forms an ionic interactions R262 within TM 6, an aromatic residue that faces into a hydrophobic core with other aromatics, and C142 which forms the last disulfide bond within the ECD. Given that the stability helix appears essential for stabilizing the ECD and the ECD is shared across the SLC22 family, we wondered whether the stability helix was also conserved. We generated a sequence alignment of human SLC22s and observed that the short motif composing the stability helix (133-SIVTEFNLVC) is highly conserved across SLC22s, in agreement with it being extremely sensitive in our mutational scan (Figure 3G). This combination of conservation and sensitivity strengthens the idea of the stability helix plays a key role in coordinating proper folding across the SLC22s.

Residues between interfaces of TMs1-6 tend to be enriched in hydrophobic residues that are sensitive to mutations implying there is a hydrophobic network that connects in the ECD to the intracellular face of the N–terminal transmembrane bundle. At the intracellular side of this bundle, TMs 2-4 and residues within the large intracellular loop between TMs 6 and 7, numerous charged residues appear to contribute to stability including D171, R175, K176, E226, and E284 (Figure 3F). In agreement with these residues playing a key role in folding and stability, these residues are extremely sensitive to mutations with charge swaps being particularly deleterious. Contrary to our expectations, these charged residues are not forming ionic interactions, pointing to the presence of intermediate conformational states with relevance to folding. While the N-terminal half of OCT1, contain most of the stabilizing residues there is a small cluster of residues in TMs 10 and 11 that are particularly sensitive to mutations, including a pair of charged residues (E455 and R462) which may form an ionic bond and are sensitive to mutations that change charge (Supp Fig 9). Interestingly, many of the charged residues in both the N and C terminal clusters of residues do not make ionic interactions in the AlphaFold2 model, suggesting additional structural states are necessary to understand folding.

A limitation of the abundance assay is that it measures overall OCT1 expression, not the surface-expressed subset. Despite this, we expect variants with high folded abundance in our assay to, in general, have high surface expression as well, barring changes to specific trafficking or targeting motifs. To validate this in OCT1, we expressed a subset of (loss of function variants and compared their total abundance and surface expression using fluorescent microscopy (Figure 3K Supp Fig 10). WT OCT1 was observed to localize to the plasma membrane. Two mutations (D149N and E386K) that are not functional based on the cytotoxicity scores but are expressed based on the abundance assay show similar surface localization to WT implying these variants disrupt substrate uptake. In contrast, three variants (W64R in the ECD hydrophobic core, V135I in the stabilizing alpha helix, and E284A in the intracellular stabilizing region) that had loss of function abundance and cytotoxicity fitness scores displayed little to no surface localization. We thus find that the mutational impacts from our high throughput screens closely follows the impact of surface expression based on microscopy. Predominantly loss of function phenotypes are due to impacts in expression and using these data we’ve determined how mechanistically these mutations are disrupting folding and stability to impact human physiology.

### A comprehensive model of structure-function of OCT1 without experimental structures

While most loss of function mutant phenotypes are due to impacts in expression, a portion of mutations minimally disrupt expression but have substantial effects on function. To understand and contextualize how mutations in OCT1 alter function, we sought to develop a structure-function model of OCT1. In SLC transporters such as OCT1, substrate uptake requires the passage through outward-facing states that can bind to a substrate and inward facing states that move a substrate inside a cell. As with all proteins, OCT1’s conformational ensemble exist within a dynamic equilibrium that is shifted by the presence or absence of substrate. The AlphaFold2 model, as a single state, is insufficient to explain how transport works because it does not inform us on what other states look like nor the interactions that drive the cycle. However, as with many transporter families, there are few experimental structures for the SLC22 superfamily, and existing structures are limited in conformational and liganded states, making it difficult to understand function within the context of the entire transport cycle. Such a comprehensive understanding of OCT1 structure-function is necessary if we want to understand how genetic variation breaks function.

We therefore adopted a new hybrid approach that combines high-quality structure prediction, coevolutionary information, MD and deep learning to reconstruct the entire functional cycle of transporters (Mitrovic *et al*., 2022). By starting from an inward-facing structure generated by AlphaFold2 we identified potential dynamic modes through a direct coupling analysis (DCA) procedure. We could then verify the existence and explore the energetics of the dynamic modes using MD simulations in conjunction with enhanced sampling. This approach reveals states that include the outward open, outward occluded, occluded, inward occluded, and inward open with and without OCT1 substrate MPP^+^ which are highly similar to the experimental solved structures in limited states (Fig 4A-C, Supp Fig 8, Supp Table 1). We included the substrate MPP^+^ because it is a canonical OCT1 cationic substrate and chemically quite similar to the substrate, SM73, we used in our cytotoxicity screen. Without the substrate, the most stable states are the outward open and occluded states. In contrast, upon binding of the cationic substrate, MPP^+^, the conformational landscape tilts towards the inward open state. This reshaping of the landscape is consistent with a cycle in which a substrate-bound OCT1 in the outward-facing state spontaneously transitions to the inward facing states, ready to release the substrate into the cytosol, and in which the empty transporter spontaneously resets to the outward facing state. As influx and efflux have been seen in OCT1 and it’s transport is not coupled to the exchange of other ions, likely the energetics of this cycle would be inverted if there are higher concentrations of substrate on the inside of the cell vs the outside (Kim *et al*., 2017).

**Figure 4.**
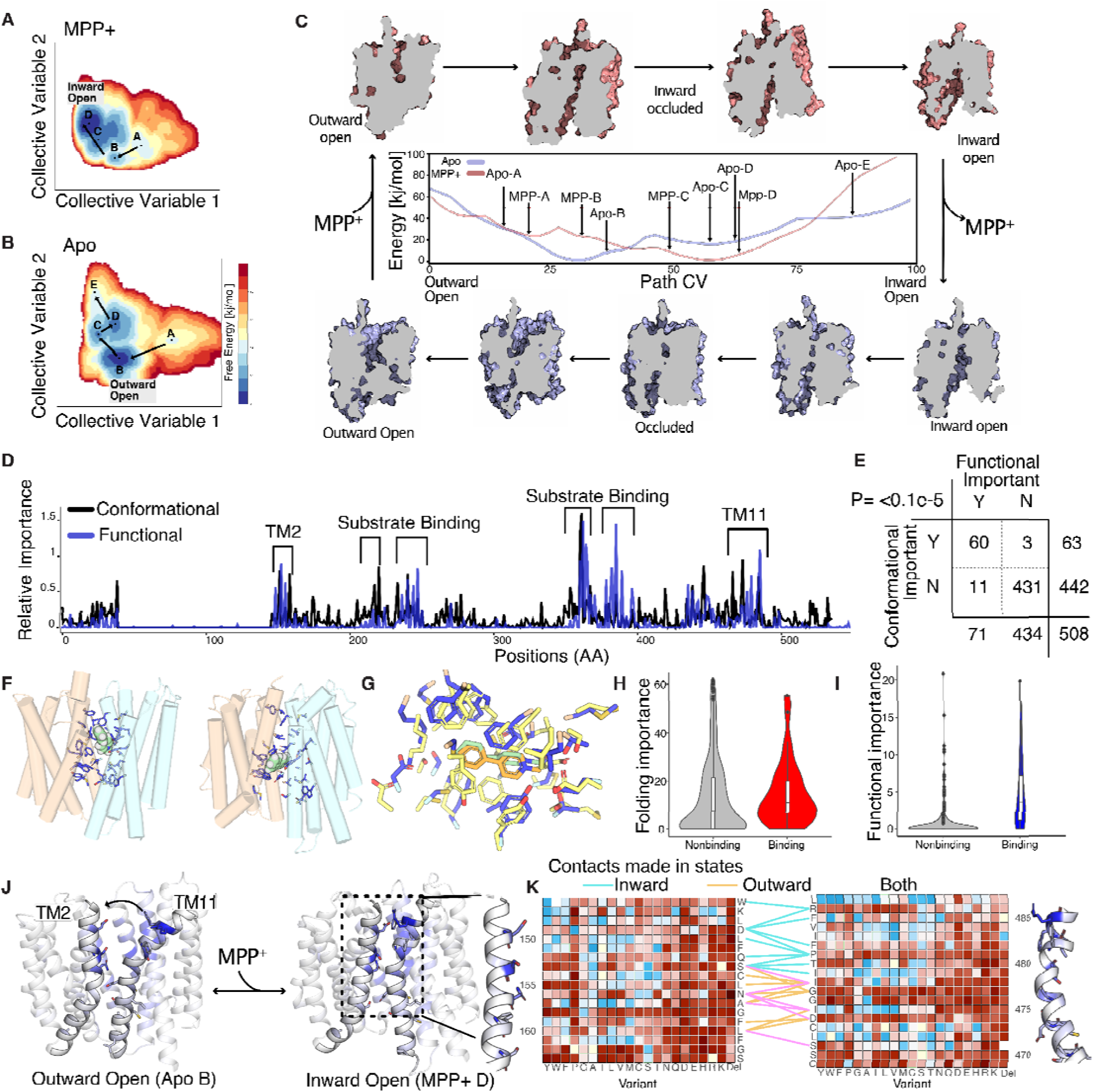
A structure-function model determined with computational structural biology and functional DMS experiment. **(A-B)** 2D conformational landscapes of apo and MPP^+^-bound OCT1 determined using MD simulations enhanced using collective variables derived from coevolutionary-based neural networks initiated from the AlphaFold2 model. The free energy landscapes are projected onto a 2-dimensional collective variable space combining contributions from residue-residue contacts specific to outward– or inward-facing states. Letters indicate the local minima in the free energy landscapes corresponding to ensembles of metastable configurations.. The paths between states are indicated with arrows leading from the outward-open to the inward-open state. **(C)** The full modeled conformational cycle of OCT1 going from MPP^+^-bound (pink) outward open (top left) to MPP^+^-bound inward open (top right), then following substrate release (purple), the inward open apo (bottom right) to outward open apo state (bottom left). In the center is a 1D projection of the energy landscapes from panels A and B, with the x-axis depicting a path collective variable representing the progression of the transporter from the outward-open to the inward-open states. The apo landscape is in light blue and MPP^+^-bound state is in light red. Indicated on the landscape are where each of the modeled states are in relation to the 1D representation in panels A,B. **(D)** DMS-based functional (blue) and MD-based conformational (black) importance scores plotted across the OCT1 sequence. **(E)** Truth table comparison across conformation and functional importance classes with a two-sided Fisher exact test showing strong significance, P Value= <0.1E-5. **(F)** Examination of substrate binding. Residues within 5 Å of the substrate (MPP^+^, in green) in any state are depicted in blue with sidechains and defined as substrate-binding residues. **(G)** The outward MPP^+^-bound state, MPP-B with residues in blue and substrate in green, overlaid with solved experimental structure, 8ET9, with residues in yellow and substrate in orange demonstrate remarkable similarities in MPP+ poses and residue placement. Residues for **(H-I)** A comparison of substrate-binding residues (as defined in **F)** with folding (**H,** in red) or functional importance (**I,** in blue) shows that MD-derived categorization and DMS-derived classification agree. **(J)** The interface between transmembrane helices 2 and 11 is highlighted here due to both their large conformational changes in the structure and enrichment of functionally important residues. Outward open and inward open states are shown with functional importance plotted and high functional importance residues modeled. **(K)** Examination of the functional heatmap for the TM2-11 interface suggests the physical logic of state-dependent interactions in OCT1. Contacts between Cα atoms within 8 Å in inward-open (cyan), outward-open (wheat), or both (magenta) states are indicated.

From the computational prediction of conformational ensembles with and without substrate, we can begin to propose structural models of substrate uptake. However, from computational structural biology alone, it is difficult to know how residues functionally contribute to conformational changes without taking all functionally relevant metastable states into consideration. To identify which residues contribute to function, we calculated a functional importance score, similar to the stability score, but now defined as the summed absolute value of cytotoxicity scores at each position across all mutations neutral for abundance (Fig 4D). To directly compare each residue’s contribution to the conformational cycle, we also quantified interaction networks between contacting residues by reweighting the entire free energy landscapes onto all inter-residue pairs, followed by calculating the betweenness centrality for each residue. This yielded a computational estimate of the residue-wise importance directly derived from the free energy landscapes while taking all conformations into consideration. We find that the computationally and experimentally determined functional scores are tremendously similar to each other allowing us to validate the mutational scanning and the predicted conformational states (Fig 4D-E).

Next, we set out to understand how functionally important residues within OCT1 contribute to function. A long standing question within the SLC22 field is whether the ECD is directly involved in function. A previous study found swapping the ECD between homologs results in changes in substrate specificity however we strongly find not a single residue essential for uptake is present within the ECD (Fig 4D, (Meyer *et al*., 2022)). Perhaps the ECD indirectly guides substrates into the substrate binding and some residue could alter binding – however as we did not include multiple substrates within the screen we did not test for this role. The role of some functionally important residues is quite clear such as those located in the substrate binding pocket, defined as being within 4 angstroms of the substrate within any of the MPP^+^ states, which likely play an important role in substrate binding and/or selectivity (Fig 4F–G). As would be expected, the substrate binding residues are extremely enriched in function-critical residues and not in those that are reported important for folding (Fig 4H–I). Within the substrate binding pockets there are several negatively charged residues that could be coordinating the movement of substrate through the pathway with the strongest function phenotype within E386. We find that negative residues mutations within the substrate binding pocket are extremely deleterious, with a particularly drastic effect of charge swaps (Supp Fig 11B). In addition, there are many aromatic residues that likely help to coordinate substrate passage through OCT1. Supporting a functional role for aromatic residues within the binding pocket, we find that all mutations to nonaromatic residues are extremely deleterious (Supp Fig 11A). In fact these residues closely resemble the binding pocket for an inhibitor that was solved in the homolog OCT3 and even a recently experimentally solved structure of OCT1 with MPP^+^ bound in the outward open state has strikingly similar placement of MPP+ between experimental and computational structures (Supp Fig 8 and 12, Khanppnavar *et al*., 2022; Suo *et al*., 2023).

Other functionally important residues make contacts within specific conformational states, implying that they play a crucial role in stabilizing those states. Amongst the most notable conformational changes from the outward to inward state is the movement of the top of TM11 towards TM2 (Fig 4J). In the inward-open facing state, TM2 and TM11 appear to make many contacts at the top of the transmembrane domains, likely stabilizing the inward-facing states (Fig 4D). In contrast, further down the TM2-TM11 interface, there are many interactions that are maintained across states, which we interpret as forming a pivot region, while others are made in the outward-facing states alone. The regions which are making interactions across the TM2-TM11 interfaces are strongly enriched for functionally important residues (Fig 4K). In particular, D149 and R486 appear to play a key role in stabilizing the inward state, P479 forms a kink that could be important in enabling the geometries necessary for these motions. Further down, a series of glycines on both TM2 (G158 and G162) and TM11 (G474-475) provide the flexibility that is necessary for the conformational changes. In addition, a key interaction F158-D474 located immediately next to the flexible glycine hinges helps to stabilize the outward state. Therefore interactions between TM11 and TM2 stabilize the transitioning through the conformational cycle. The conformational ensembles complemented by multi-phenotypic mutational scanning data generated here thus allow us to propose a structure-function model for OCT1 in absence of experimentally solved structures.

### Inferring the biophysical mechanisms by which mutations alter OCT1 function

By understanding how OCT1 likely functions, we can contextualize how variants could break OCT1 function in people. We next sought to develop a biophysics-based explanation for how mutations alter function and focused on a subset of 20 mutations which span loss of function and gain of function phenotypes, are within functionally or stability important regions, and are when possible, predominantly human polymorphisms (x are human). To ensure that variants are indeed loss of function, we conducted lower-throughput radioligand uptake experiments in HEK293T cells for all 20 with a range of substrates including two canonical canionic OCT1 substrates MPP+ and TEA+ as well as pharmaceuticals sumatriptan and metformin (Fig 5A-B, Supp Figs 13). We find that radioligand uptake experiments recapitulate the impacts within the screens. Unsurprisingly, MPP+ which is chemically most similar to SM73 has the highest correlation between radioligand uptake experiments and the cytotoxicity screen.In some cases variants which are known to be loss of function clinically weren’t in our screen, such as a common variant M408V. For example, in our validation experiments we found variants (R486E and R488E) which were not loss of function in the screen also were not with MPP+ but were strong loss of function with clinically relevant substrates, Metformin and Sumtriptan (Supp Fig 13). This implies they impact substrate specificity, which is known to be dependent on polymorphisms. In the future, it will be important to screen transporters across a range of substrates to understand better how different variants tune substrate specificity and perhaps also reveal the mechanistic basis of polyspecificity.

**Figure 5.**
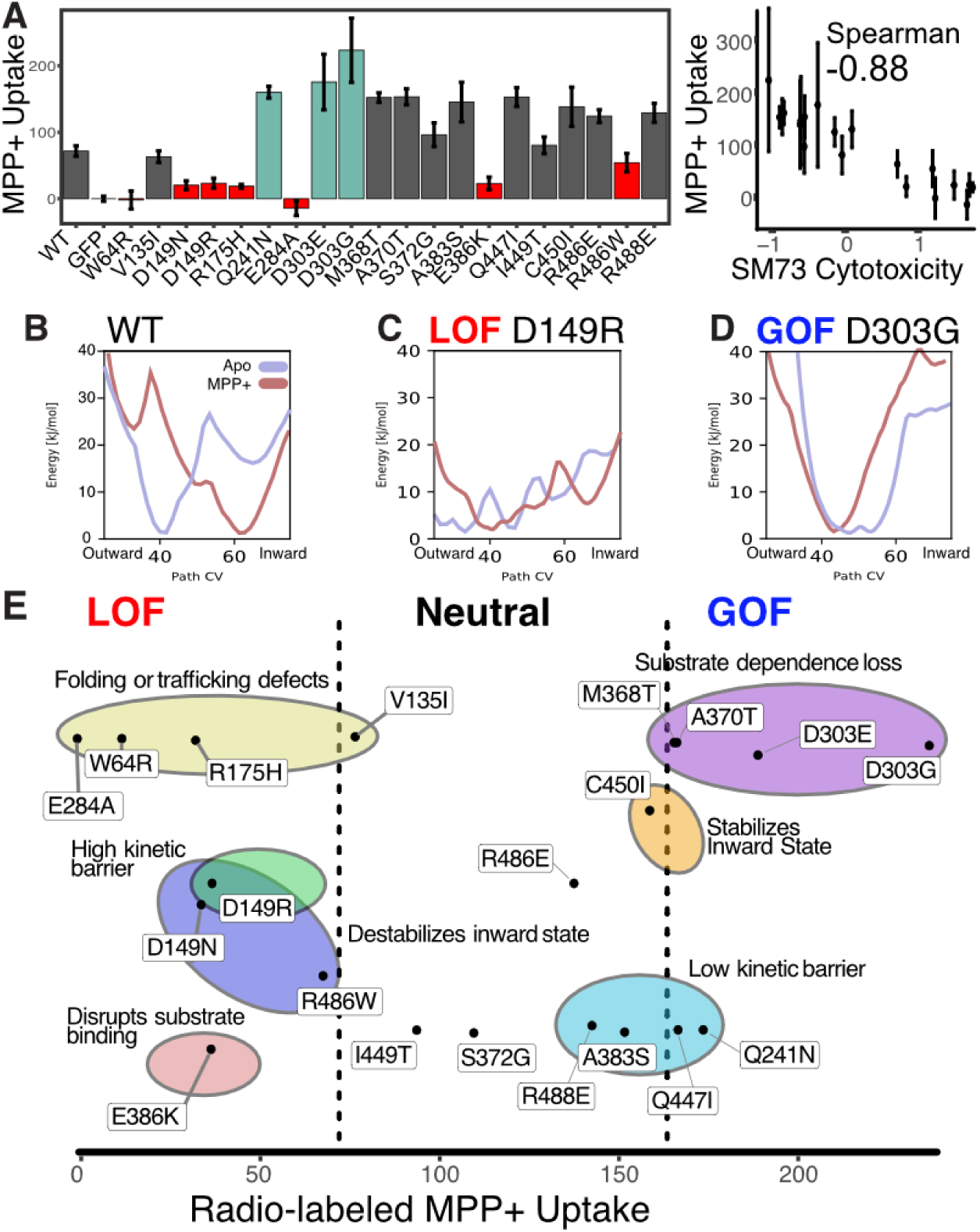
Biophysical basis of mutations on OCT1 function. **(A)** Radio-labeled MPP^+^ uptake experiments for a subset of mutations across a range of fitness mechanisms and effects. Left, mean experimental uptake across 3 replicates (mean with error bars, SEM). Right, correlation between SM73 cytotoxicity fitness scores and radio-labeled MPP^+^ uptake scores (error bars, SEM), with rank-order Spearman correlation coefficient shown. **(B-D)** 1D free energy landscapes of the rocker-switch motion for WT, loss of function mutant D149R, and gain of function mutant D303G. Apo landscapes are in light blue and MPP^+^-bound in light red. **(E)** Mechanistic basis of mutational impact on OCT1. Mutations are plotted based on their MPP^+^ uptake scores and classified based on loss of function or gain of function using the high-throughput screen results. Mechanistic classifications inferred from their respective free-energy landscapes (Methods) are grouped within colored circles.

Some of the mechanistic impacts can be directly intuited such as the effect of mutating the substrate binding residue E386 to lysine: introducing a positively charged residue will likely lead to diminished substrate binding and thus loss of transporter function. Others that can be immediately intuited are mutations within residues with high stability importance (W64R, V135I, R175H, E284A), which are likely disrupting folding. In other cases where mutations which alter substrate uptake are likely changing the conformational ensemble, it is more challenging to immediately intuit the effects of a variant.

To develop a mechanistic understanding for mutations within OCT1, we conducted the same computational structural biology pipeline we used for getting structural states and predicted the conformational landscape with and without the substrate MPP^+^ for all mutants (Fig 5B-D, Supp Fig 14-15). Using fitness scores, conformational ensembles, and comparing the thermodynamic stability of inward– and outward-facing states, we developed a biophysics-based understanding for how each mutation disrupts OCT1. For example, we classified mutations within the hydrophobic core, which we propose drive loss-of-function as folding defects (Fig 3K, Supp Fig 10). From the free energy landscapes derived from the MD simulations, we find examples in which mutations alter the conformational ensembles in a manner that can lead to loss-of-function: for example, D149R and D149N at the TM2-TM11 interfaces destabilize the substrate-dependent exchange between outward and inward states by increasing the energetic barrier – an effect we name high kinetic barrier (Fig 5C, Supp Fig 14-15). In addition, D149R is no longer stabilized in the inward-facing state in the presence of bound MPP+, and we conclude that loss-of-function also stems from a transporter arrested in the outward-facing state (an effect we name destabilizes the inward-facing state). We were also interested to discover how gain-of– function mutations impacted OCT1. For example, D303G changes the ensemble such that the energetic barriers between outward and inward states are dramatically reduced and barely change between substrate addition. Effectively, this will increase the exchange between outward– and inward-facing states and increase the rate of substrate transport into the cell. A similar mechanism is proposed to have evolved in the malarial sugar transporter PfHT1 where stabilization of the transition state enables promiscuous cycling of a wide range of substrates with high turnover (Qureshi *et al*., 2020). Combining mutational scanning and computational ensemble predictions allows us to propose structure-function models for how OCT1 moves substrates into cells. Furthermore, we show we can predict how human polymorphisms change the energetics of transporter function which tells us mechanistically how an important pharmacogene’s variation changes a transporter.

### An atlas of OCT1 human variant impacts and mechanisms of action across populations

To understand how OCT1 variation ranges across people and populations, we compared our mechanistic mutational scanning data to a large-scale public database that provides a comprehensive and diverse collection of human genetic variation, gnomAD (Genome Aggregation Database). gnomAD v2.1.1 contains 160 synonymous, 392 missense, and 3 in-frame deletion and 2 in-frame insertion variants (Karczewski *et al*., 2020). Prior to this work, only 39 of these variants had been characterized, whereas now due to our DMS experiment nearly all variants in people have functional annotation (Fig 6A-B). Between the 23 that had been previously studied and are present in our DMS, we find a strong concordance with published MPP^+^ uptake experiments in the literature with the cytotoxicity (p<0.0001, Pearson r = –0.9) and abundance (p<0.0001, Pearson r = 0.91) experiments (SI Table S3). After observing 375 missense variants and 2 in-frame deletion human variants, we found that 91 out of 368 (25%) of them had significant loss of function phenotypes based on cytotoxicity, while 125 out of 377 (34%) had significant loss of protein abundance (Fig 6B-C). Within the 86 human loss of function variants folding is the primary driver with only 10 variants having disrupted substrate uptake. This is consistent with previous studies of membrane proteins in which folding is the primary determinant of loss of function including the homolog OCTN2 (Koleske *et al*., 2022). While folding primarily explains loss of function overall, there are numerous variants with reduced expression yet like wildtype substrate uptake which implies that while expression is decreased there is sufficient OCT1 at the surface to uptake the cytotoxic substrate. This could imply a differential sensitivity between the abundance and cytotoxicity assays which could be due to overexpressing OCT1 during screens. Perhaps if we tested multiple levels of substrate or reduced expression we could detect a substrate uptake effect of these mild loss of function variants.

**Figure 6.**
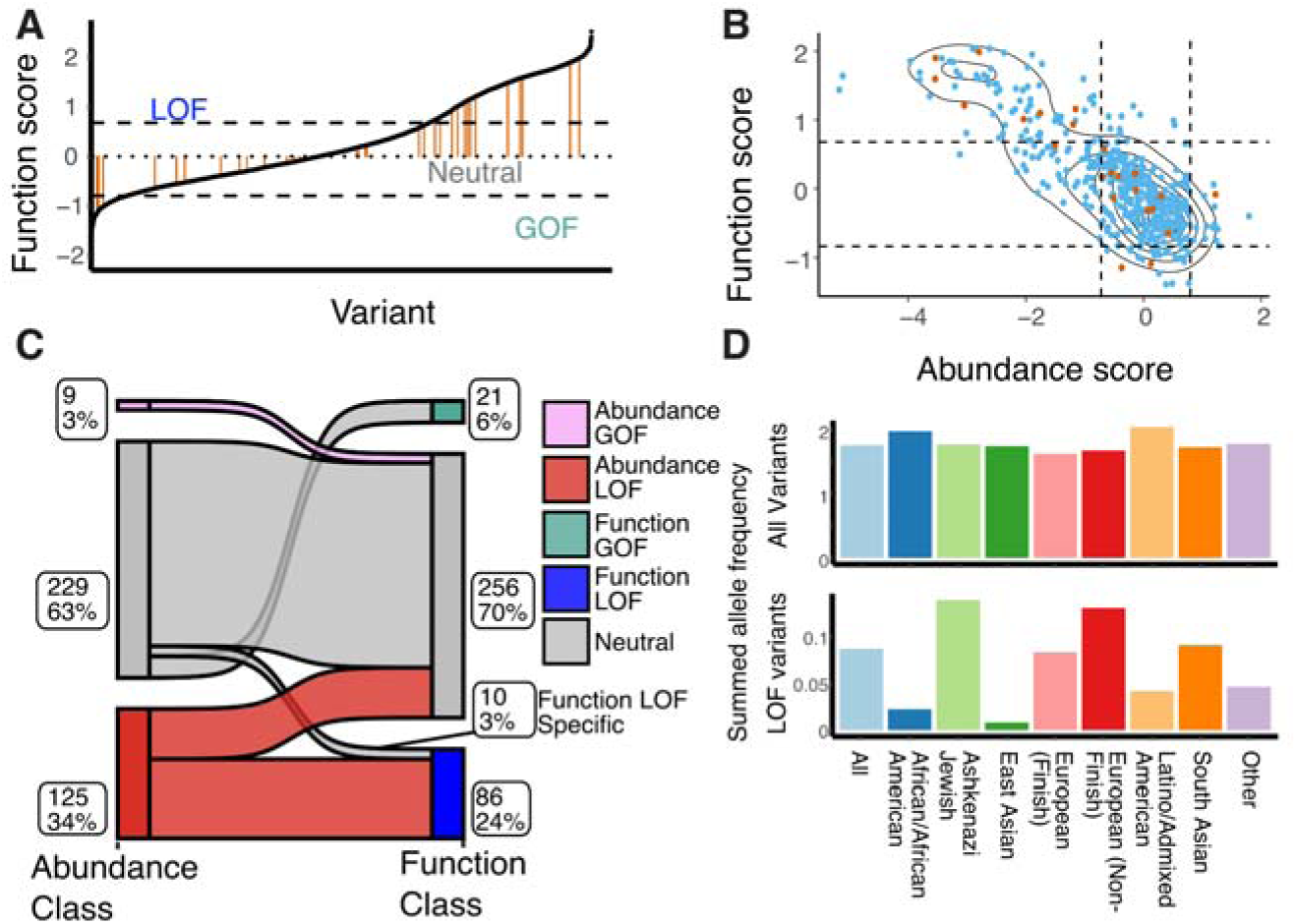
Mechanism of action of OCT1 polymorphisms. **(A)** Distribution of previously characterized human polymorphisms from gnomAD (39 variants, orange) among the total set from this work is shown among the rank-ordered set of all 515 gnomAD variants. Cutoffs are shown based on the synonymous mutation distributions as dotted lines. **(B)** Abundance and function scores plotted for all human variants, showing previously-studied variants (orange) among all previously uncharacterized (blue). Cutoffs are shown based on synonymous mutation distribution as dotted lines. **(C)** Classification of human variants by their predicted impact based on synonymous-derived cutoffs. The diagram shows how observed functional variation (right) is conditioned on abundance impacts (left). Numbers and percentages of variants within each group are next to the respective subpopulation. Flows between classes suggest the interaction between abundance and function. **(D)** Summed allele frequencies across populations with differing ancestry within gnomAD reveal the utility of functional annotation for understanding human variation. Across all variants, all groups have similar alternative allele frequencies (top), but looking at only loss of function variants reveals large differences (bottom).

For human variants, we sought to use the mechanistic understanding of folding and function we determined to contextualize how OCT1 breaks. We found that many residues, primarily within the N-transmembrane domain, contribute to folding and so it should come as no surprise that folding disrupting human variants are similarly widely distributed. This includes mutants we validated in microscopy and uptake experiments within the ECD hydrophobic core (W64R), disulfide forming cysteines (C88R), the stability helix (V135I), and residues making ionic interactions within the intracellular adjacent helical bundles (R175H). While folding primarily disrupts OCT1 function within human variants, we also developed an intuition for several of the 10 variants which specifically impact substrate uptake. Some human variants that disrupt substrate uptake impact substrate binding directly, such as E386K and R439Q. In other cases mutations likely disrupt the flexibility of key flexible or rigid regions necessary for conformational changes such as G158A within the TM2 which moves to interact with TM11 as described above or P388S which is at a helix break near the substrate binding E386. Other function specific residues even destabilize the TM2-11 interactions directly such as (F485V and R486W). Biophysically, loss of function with OCT1 spans the wide-ranging interactions that drive stability, ligand binding, and conformational exchange that is necessary for substrate uptake.

With the 39 human variants which were previously characterized, it is difficult to determine whether there is a depletion of loss of function variants in OCT1 across populations. We explored whether reduced function variants were evenly distributed across all populations. Surprisingly, when comparing the six loss of function variants with mean allele frequency (MAF) ≥1%, we found that they were absent in East Asians, with MAF <0.0005%. In fact, East Asians had the lowest total allele frequency of common poor functioning variants at 0.1%, followed by African (7.4%), South Asian (14.8%), European (20%), Non-Finnish European (27.4%), Ashkenazi (26.1%), and Latino (26.2%) populations, all of which had much higher total allele frequencies. Furthermore, when considering all common and rare variants, we identified 141 poor function across all populations. Consistent with our earlier observation, East Asians had the lowest total allele frequency at 1.1%, followed by African (7.8%), South Asian (16.1%), European (20.1%), Non-Finnish European (27.9%), Ashkenazi (26.5%), and Latino (26.5%) populations, all of which had much higher total allele frequencies (Fig 6D). A previous study observed the strongest divergence in the frequency of losing OCT1 activity between populations in America and East Asia (Seitz et al., 2015). Without a systematic functional study like this deep mutational scan, it is challenging to demonstrate global variations in the frequency of loss of OCT1 activity

### Poor functioning OCT1 variants broadly result in increased cholesterol levels in humans

We next sought to understand how OCT1 variation contributes to the diversity of human physiology. Poor function OCT1 polymorphisms are linked to increased LDL cholesterol, total cholesterol, and triglycerides (Liang *et al*., 2018). Therefore, we hypothesized that poor functioning variants from our screens should also have a commensurate increase in these markers. To compare our unbiased mutational scan to human physiology, we relied on the UK Biobank (UKB), which contains exome sequencing and associated patient health records. Within OCT1, the UKB database contains 268 missense variants and 1 deletion variant of which 264 variants are in the OCT1 DMS (SI Table S5-6). Of the 12 variants with a MAF >0.1% in the UKB population, six have poor OCT1 function either through our experiments or from previous literature (R61C, C88R, S189L, G401S, M420del, and G465R). Within all of these mutations individually and together we find a strong association with increased LDL, cholesterol, and triglycerides (SI Table S6-7). We used our DMS data to assign loss of function parameters to all missense variants and in-frame deletion within the UKB. We find that poor functioning SNPs within the 180,000 European genomes present within the dataset are significantly associated with LDL (p=2.8×10^-10^), total cholesterol (p=1.1×10^-9^), and triglyceride (p=5.8×10^-11^) levels (SI Table S8). In contrast, the 142 SNPs which are similar to wildtype have far weaker associations with LDL (p=0.04), total cholesterol (p=0.088), and triglyceride (p=0.02) levels showing weaker associations. In contrast, comparing other molecules such as uric acid which are not OCT1 substrates, we find no associations at all (SI Table S8). These results imply that DMS experiments such as OCT1 can infer the impact of genetic variation within transporters on human physiology.

## Discussion

By measuring how mutations alter OCT1 abundance and substrate uptake, we discover that loss of function mutations are overwhelmingly caused by disruptions in expression and folding. This result follows observations we have made previously in a deep mutational scan of a potassium channel, Kir2.1, and a more limited mutational screen of human variants within a Carnitine transporter, OCTN2 (Zhang *et al*., 2021; Coyote-Maestas *et al*., 2022; Koleske *et al*., 2022). Other membrane protein-associated diseases are also caused by folding defects, such as in the chloride channel CFTR and GPCR Rhodopsin (Lukacs and Verkman, 2012; Penn *et al*., 2020). Approaches such as VAMP-seq which measures protein abundance provide a powerful and generalizable technique to study membrane protein folding. Using this expression screen, we identified the stability helix and ECD domain as a key determinant folding across SLC22 transporters with little prior studies. Surprisingly the N terminal 6 TMDs of OCT1 primarily contribute to folding while the C terminal 6 TMDs do not. As the second 6 TMDs are more enriched for functionally critical residues, this implies a hierarchical organization in which some residues drive stability whereas others underlie function. By applying these approaches more broadly in other MFS transporters, we could determine whether this is a general phenomenon, which could explain the benefit evolutionarily of the fusion of two 6 TMD domains which could provide a ‘division of labor’. Such broad datasets across the membrane proteome could yield biophysically informed generalizable machine-learning models to understand how membrane proteins fold and how mutations break them. The causes of defects in membrane protein folding have been difficult to understand due to immense technical challenges; however, here we show that mechanistic genetic screening provides a long-sought avenue to develop mechanistic models. Such models would provide fundamental insights into how membrane proteins fold and could guide the development of therapeutics that stabilize the disruption of folding by mutations such as the blockbuster CFTR chaperones (Fiedorczuk and Chen, 2022; McKee *et al*., 2022).

We develop mechanistic models for how OCT1 works and how mutations alter conformational ensembles by integrating our high throughput functional screen with computational structural modeling. Recent breakthroughs in experimental and computational structural biology allowed the membrane protein field to begin to understand more broadly how membrane proteins function (Cheng, 2018; Del Alamo *et al*., 2022). An immensely powerful resource to aid in this is the development of approaches which can predict entire conformational landscapes as highlighted in this work (Fleetwood *et al*., 2020; Fleetwood, Carlsson and Delemotte, 2021; Mitrovic *et al*., 2022). By integrating structural computational biology and mechanistic mutational scanning, we can begin to understand the complete conformational landscape of cross-membrane transport and how human variants alter these landscapes. Mutational scans could identify mutations which trap structures in specific states to test hypotheses generated by computational structural biology and resolve unstable states. Perhaps even large-scale thermodynamic double mutational cycles could be used to refine conformational states and ensembles similar to how double mutational scans could reveal the structure of a small protein (Olson, Wu and Sun, 2014; Rollins *et al*., 2019; Schmiedel and Lehner, 2019).

Amongst the most challenging problems for transporter biology is understanding substrate specificity. In pharmacogenomics, we currently cannot predict why or how human polymorphisms will differentially impact drug uptake. For structural biology, this problem is intractable as it’s not feasible to solve structures of say OCT1 and its homologs with the hundreds of known substrates (Gebauer *et al*., 2022). A major limitation of this study is that we only used a singular substrate for our screen– as it stands with these data, we also cannot fully understand the chemical-genetic interactions that give rise to the diversity of drug responses in people. In the future, mutational scans complemented with computational modeling could enable a far more rigorous and systematic approach. At a first level this sort of data would aid in understanding and predicting the differential impacts of mutations on specific substrates in people for precision medicine. Perhaps if we develop a mechanistic understanding of residue-chemical interactions we could better understand the residue-chemical interactions that guide substrate specificity we could engineer transporters specific for a substrate, design more substrates with reduced side effects or tissue specificity, or if our models become sufficiently sophisticated – de-orphan transporters.

Pharmacogenomic research on OCT1 unveiled population specific variants, yet without systematic studies of variant effects it is infeasible to study the distribution of functional variants between populations (Shu *et al*., 2003; Seitz *et al*., 2015). Though OCT1 genetic polymorphisms are not yet routinely monitored in clinical pharmacogenetic analyses, there is a wealth of data suggesting that combining poor function polymorphisms would lead to significant differences in drug levels and response, for examples metformin (Shu *et al*., 2007), fenoterol (Tzvetkov *et al*., 2018), tramadol (Stamer *et al*., 2016), morphine (Tzvetkov *et al*., 2013) and sumatriptan (Matthaei *et al*., 2016). Further, as sequencing is increasingly adopted for clinical pharmacogenomic testing (as opposed to genotyping), it will be essential to understand the function of OCT1 mutations uncovered during routine pharmacogenomic testing. In this study, we demonstrate the feasibility of combining data from UKBiobank and DMS to investigate the effects of normal and loss of function variants in OCT1 on cholesterol levels, a known phenotype associated with OCT1 polymorphisms. Future studies are required to further investigate the potential translation of this information to drug response and toxicity. By integrating our DMS results with gnomAD, a comprehensive collection of population variations, to infer variant effects in populations and individuals, we have made an intriguing observation. Overall, we find that common missense mutations in humans generally retain normal function with the loss of function human variants most commonly due to folding defects. Individuals of East Asian and African ancestry demonstrate a surprising lower prevalence of loss of function polymorphisms. There are several possible explanations for differences in the frequencies of reduced function alleles among populations, ranging from selective evolutionary pressure for maintaining or losing OCT1 function within specific populations to random bottlenecks. It will be exciting to identify the specific mechanism that accounts for the differences in functionality across populations. For example, previously, we observed that loss of function in OCT1 could confer a selectable advantage, as evidenced by the protective phenotypes against thiamine deficiency observed in Oct1 knockout mice (Liang *et al*., 2018). The distinct functionality of OCT1 across different populations can have profound implications for clinical outcomes. For instance, morphine has been observed to exhibit stronger efficacy in East Asian populations, requiring significantly lower doses compared to Europeans (Konstantatos *et al*., 2012; Letchuman *et al*., 2023). Further investigation is needed to examine the impact of polymorphisms in morphine metabolizing enzymes and transporters. An unbiased understanding for how variants differentially impact drug transporters within populations would allow for personalized clinical decision-making, customized dosage, and improved treatment.

Broadly applied, such approaches such as those applied in this study would yield a biophysics-based understanding of evolution, a deep understanding of protein biology, human genetic variation, and the diversity of life. The integration of mechanistic mutational scanning, biophysical models of folding and structure, and large scale patient databases would allow a mechanistic prediction for the impact of variation. This would transform our basic biology for how proteins underlie our physiology, yet also guide the diagnosis of disease, and development of therapeutics to better treat disease.

## Methods

### Library generation

The OCT1 library generation was performed using the previously described method, DIMPLE (Deep Insertion, Deletion, and Missense Mutation Libraries for Exploring Protein Variation in Evolution, Disease, and Biology) (Macdonald *et al*., 2023). This method was adapted from SPINE and was used to design oligo to create mutations, insertions, and deletions (Nedrud, Coyote-Maestas and Schmidt, 2021). The code was used to allow one to mutate each amino acid position to the other 19 amino acids, mutate to a synonymous codon, and a deletion (see code deposited at https://github.com/coywil26/DIMPLE). Primers and oligos that are designed and used for generating OCT1 libraries are listed in Supp Table 9-10.

### Library generation and cloning

Method for library generation and cloning was described previously (Macdonald *et al*., 2022). Oligos for SLC22A1 was synthesized by Agilent (SurePrint Oligonucleotide, Agilent Technologies) to give 10 pmol of lyophilized DNA, which was then resuspended in 1 X TE buffer. Vector containing SLC22A1 was synthesized by Twist Bioscience in the High Copy Number Kanamycin backbone, the lyophilized plasmid DNA was resuspended to 10 ng/µL in 1 x TE buffer.

Sublibraries of different regions of SLC22A1 were PCR amplified using primer-specific and polymerase (PrimeStar GXL DNA polymerase) (Takara Bio). A total of 11 regions were PCR amplified, each in 50 µL reactions using 1 µL of the total OLS library as template and 30 cycles of PCR. 5 µL of the reactions was assessed by running on an agarose gel. The reactions were cleaned up using Zymo Clean and Concentrate kits (Zymo Research) and eluted in 10 µL of TE buffer. For each 11 region, the plasmid was amplified to add on golden gate compatible Type IIS restriction sites complementary to those encoded within the sublibrary oligos using Primestar GXL polymerase according to the manufacturer’s instructions in 50 µL reactions using 1 µL of the template vector and 30 cycles of PCR. The entire PCR reaction was run on a 1% agarose gel and gel purified using a Zymoclean Gel DNA Recovery Kit.

Target gene backbone PCR product and the corresponding oligo sublibrary were assembled using BsaI-mediated Golden Gate cloning. Each 40 µL reaction was composed of 300 ng of backbone DNA, 50 ng of oligo sublibrary DNA, 2 µL BsaI-HF v2 Golden Gate enzyme mixture (New England Biolabs), 4 µL 10× T4 Ligase buffer, and brought up to a total volume of 40 µL with nuclease free water. These reactions were placed in a thermocycler with the following program: (i) 5 min at 37°C, (ii) 5min at 16°C, (iii) repeat (i) and (ii) 29 times, (iv) 5 minutes at 60°C, (v) hold at 10°C. Reactions were cleaned using Zymo Clean and Concentrate kits, eluted into 10 µL nucleus free water, and transformed into MegaX DH10B electrocompetent cells (Thermo Fisher) according to manufacturer’s instructions.

MegaX DH10B cells were recovered for one hour at 37°C. A small subset of the transformed cells were plated at varying cell density to assess transformation efficiency. All transformations had at least 100× the number of transformed colonies compared to the library size. The remaining cell outgrowth was added to 30 mL LB with 50 µg/mL kanamycin and grown at 37°C with shaking until the OD reached 0.6 – 0.7. Library DNA was isolated by miniprep (Zymo Research). Sublibrary concentration was assessed using Qubit. Each sub-library of a given gene was pooled together at an equimolar ratio. These mixed libraries were assembled with a landing pad cell line compatible backbone containing a Carbenicillin resistance cassette and GSGSGS-mNeonGreen Fragment P2A-Puromycin cassette for positive selection.

Libraries were cloned into a landing pad vector containing a BxB1-compatible a*ttB* recombination site using BsmBI mediated golden gate cloning. We kept track of transformation efficiency to maintain library diversity that was at least 100× the size of a given library. We designed the landing pad vector which we recombined the library into to contain BsmBI cutsites with compatible overhangs for the library to have an N terminal kozak sequence and in-frame with a C-terminal GSGSGS-mNeonGreen Fragment P2A-Puromycin cassette for positive selection. The golden gate protocol we used was 42°C for 5 minutes then 16°C for 5 minutes repeated for 29 cycles followed by 60°C for 5 minutes before being stored at 4°C. This landing pad backbone was generated using Q5 site-directed mutagenesis, according to the manufacturer’s suggestions(Macdonald *et al*., 2023). Reactions were cleaned using Zymo Clean and Concentrate kits, eluted into 10 µL nucleus free water, and transformed into MegaX DH10B electrocompetent cells (Thermo Fisher) according to manufacturer’s instructions. MegaX DH10B cells were recovered for one hour at 37°C. A small subset of the transformed cells were plated at varying cell density to assess transformation efficiency. All transformations had at least 100× the number of transformed colonies compared to the library size. The remaining of the cells were plated into two large square plates (245mm x 245mm) containing 200 mL LB Amp.

### Cell line generation and cell culture

The cell lines generated for this study were previously described (Matreyek *et al*., 2020; Coyote-Maestas *et al*., 2022). To make the cell lines, 1500 ng of library landing pad constructs (described in previous section) were co-transfected with 1500ng of a BxB1 expression construct (pCAG-NLS-BxB1) using 10.5µL of lipofectamine LTX in 10 wells of a 6 well plate. All cells were cultured in 1X DMEM, 10% FBS, 1% sodium pyruvate, and 1% penicillin/streptomycin (D10). The HEK293T based cell line has a tetracycline induction cassette upstream of a BxB1 recombination site and split rapamycin analog inducible dimerizable Casp-9. Two days following transfection, expression of integrated genes or iCasp-9 selection system is induced by the addition of doxycycline (2 µg/µL, Sigma-Aldrich) to D10 media. Two days after induction with doxycycline, AP1903 is added (10nM, MedChemExpress) to cause dimerization of Casp9. Successful recombination shifts iCasp-9 out of frame, so only non-recombined cells will die from iCasp-9 induced apoptosis following the addition of AP1903. After two days of AP1903-Casp9 selection the media is changed back to D10 with doxycycline and cells are allowed to recover for two days. After allowing cells to recover for two days, media was changed to D10 with doxycycline and puromycin (2 µg/ml, Life Technologies Corporation), as an additional selection step to remove non-recombined cells. Cells remained in D10 plus doxycycline and puromycin for at least two days until cells stopped dying. Following puromycin treatment cells are detached, mixed, and seeded on two T75 flasks. Cells were then allowed to grow until they reached near confluence, then frozen in aliquots in a cryoprotectant media (2X HyClone, HyCryo, Cryopreservation Reagent).

### Sequencing library preparation and genomic DNA extraction and data analysis

Genomic DNA was extracted using a Quick-DNA^TM^ Microprep Plus Kit (Zymo Research) from cells sorted into four different GFP intensities. Whereas, genomic DNA was extracted using a Quick-DTNA^TM^ Miniprep Plus Kit (Zymo Research) from cells treated with or without SM73 (1 µM) at different timepoints. The extracted genomic DNA from the miniprepped or micro kit prepped plasmid library were amplified using Landing_pad_backbone_for and P2A_cell_line_rev primers (Supp Table 9).

Amplicons were prepared for sequencing using the Nextera XT DNA Library kit from Illumina with 1 ng of DNA input. Samples were indexed using the IDT for Illumina Nextera DNA Unique Dual Indexes Set D (96 Indexes) and SPRISelect beads (Beckman Coulter) at a 0.9× ratio were used for cleanup and final size selection. Each indexed tagmented library was quantified with Qubit HS as well as Agilent 2200 TapeStation. Samples were then pooled and sequenced on a NovaSeq 6000 SP300 flowcell in paired-end mode, generating fastq files for each sample after demultiplexing. Each fastq was then processed in parallel using the following workflow: adapter sequences and contaminants were removed using BBDuk, then paired reads were error corrected with BBMerge and then mapped to the reference sequence using BBMap with 15-mers (all from BBTools (Bushnell, 2014)). Variants in the mapped SAM file were called using the AnalyzeSaturationMutagenesis tool in GATK v4 (Van der Auwera and O’Connor, 2020). The output of this tool is a CSV containing the genotype of each distinct variant as well as the total number of reads. This was then further processed using a python script, which filtered out sequences that were not part of the designed variants, then formatted input files for Enrich2 (Rubin *et al*., 2017). Enrichment scores were calculated from the collected processed files using weighted least squares and normalized using wild-type sequences. The final scores were then processed and plotted using R. Read counts are reported within Supplemental Table 11. Enrich2 scores for all replicates and overall are reported in Supplemental Table 12 and 13, respectively. Our libraries and cell lines show good coverage and low bias, as desired, with >97% of variants identified in our pre-selection samples and even representation across variants and positions (Supp fig 1).

### Transient transfection of the complementary remainder of the green fluorescent protein (mNG2-1-10)

HEK293T cells with the OCT1 library of variants are transfected with the green fluorescent protein (mNG2-1-10). 600,000 – 700,000 cells/well (6-well poly-d-lysine coated) were seeded the day before the transfection. A total of 12 wells were plated and including a well containing HEK293 cells without the landing pad integration site. The mNG2-1-10 constructs (3000 ng/well) were mixed with OptiMEM media (0.5 mL/well) (Life Technologies) and Lipofectamine LTX (10.5 µL/well). Samples were mixed by pipetting up and down several times, allowed to stand at room temperature for 15-20 min. The media with the cells seeded the day before were removed and fresh media (2 mL, DMEM-H21, 10% FBS) were added to each well. After 15-20 min incubation time of the mixture above, then add 0.5 mL of the mixture to each well of the 6-well plate. After 24 hours, cell media were removed, and fresh media were added to each well. (DMEM-H21, 10% FBS). After additional 24 h, cells are ready for subsequent experiments with fluorescent-activated cell sorting (FACS) to sort the cells into different GFP abundance.

### Fluorescence-activated cell sorting for abundance assay

In VAMP-seq, we add a small fragment of a split-fluorescent protein, to the cytoplasmic C-terminus of OCT1 (OCT1-mNG2-11) than cytoplasmically co-express the complementary remainder of the fluorescent protein (mNG2-1-10). Folded OCT1-mNG2-11 efficiently assembles with mNG2-1-10 to generate fluorescence, allowing fluorescence to serve as a FACS-seq-compatible proxy for folding and stability. After conducting a 24–hour transient transfection of mNG2-1-10 (as described in the previous section), all cells were sorted using a BD FACSAria II cell sorter. mNeongreen and mCherry fluorescence was excited with a 488 nm laser and recorded with a 530/30 nm band pass filter and 561 nm laser and 585/42 nm band pass filter and gated based once mNeonGreen fluorescence doesn’t depend on mCherry fluoresence Cells expressing surface markers were sorted into four subpopulations based on their mNeonGreen fluorescence levels. As the fluorescence distribution showed a skewed pattern, we separated the gates into even bins (Supp Fig 2). Subsequently, each cell population underwent the extraction of genomic DNA and the preparation of a library for sequencing, as described in the previous method section.

### Site-directed mutagenesis to create OCT1 variants

Twenty-six missense variants were selected to validate the OCT1 function and abundance. Six of the variants (R61C, C88R, P117L, G401S, M420Del, G465) in the landing pad vector backbone generated using site-directed mutagenesis. To create each mutation, NEBaseChanger tool was used to design primers and Q5^®^ Site-Directed Mutagenesis Kit was used to perform mutagenesis. For the other twenty of the OCT1 variants, we used the site-directed mutagenesis service offered by the GenScript (New Jersey, USA). These twenty OCT1 variants are in the pcDNA3.1(+)-GFP (C-terminal). For all the constructs, Sanger Sequencing was used to confirm sequence.

### Transient transfection of plasmid containing OCT1 variants

Twenty-six missense variants were selected to validate the OCT1 function and abundance. Six of the variants (R61C, C88R, P117L, G401S, M420Del, G465) and reference OCT1 were transfected into HEK293T cells containing the landing pad and were used to create stable cell lines. The six OCT1 variants are in the landing pad vector backbone. Twenty of the variants and reference OCT1 were transfected into HEK293 cells (UCSF Cell Culture Facility) to create transient expressing cell lines. The twenty OCT1 variants are in the pcDNA3.1(+)-GFP (C-terminal). Transient transfection of constructs encoding OCT1 reference and OCT1 variants was achieved by reverse transfection using Lipofectamine LTX transfection reagent (Thermo Fisher Scientific). 50,000 cells/well (96-well) were used in the reverse transfection. Constructs (100 ng/well) were mixed with OptiMEM media (20 µL/well) (Life Technologies) and Lipofectamine LTX (0.2 µL/well). Samples were mixed by pipetting up and down several times, allowed to stand at room temperature for 15-20 min, and then added to each well of the 96-well plate (20 µL) (poly-d-lysine coated). HEK293 cells were counted and seeded into wells at a density of 45,000 cells/well (100 µL/well) (96-well). After 24 hours, cell media were removed, and fresh media were added to each well. (DMEM-H21, 10% FBS). After additional 24 h, cells are ready for subsequent experiments (uptake assays). To investigate the effects of missense variants on OCT1 expression, stable HEK293 cells expressing the variants or reference OCT1 were created. Specifically, eight missense variants (W64R, V135I, D149N, D149R, R175H, E284A, E386K, R486W) and a reference OCT1 sequence in the pcDNA3.1(+)-GFP (C-terminal) expression vector were transfected into HEK293 cells to generate stable cell lines. We subsequently analyzed OCT1 expression in these cells using confocal imaging.

### Fluorescence microscopy

For the immunostaining experiments, HEK293 stable cell lines were plated onto poly-D-lysine treated 12-well plates with sterile coverslips at a density of 200,000 cells per well. Two days post-seeding, when cells reached 90-100% confluency, they were stained. On the day of staining, the cell media was removed and cells were washed with cold Hank’s Balanced Salt Solution (HBSS, Thermo Fisher Scientific Inc.). The plasma membrane was stained first with Wheat Germ Agglutin (WGA) Alexa Fluor 647 conjugate (Invitrogen Life Sciences Corporation), diluted in HBSS at 1:500, for 15 minutes at room temperature (RT). After staining, the solution was removed and cells were washed three times with HBSS. Cells were fixed with 3.7% formaldehyde in HBSS for 20 minutes, and after aspiration, cells were washed again three times with HBSS. The nucleus was then stained with Hoechst solution (Thermo Fisher Scientific Inc.), diluted at 1:2000 in HBSS, for 20 minutes at RT in darkness. After staining, the solution was aspirated and cells were washed twice with HBSS. Coverslips were carefully mounted on Superfrost Plus Microscope Slides (Thermo Fisher Scientific Inc.) with a drop of SlowFade Gold Antifade mountant (Thermo Fisher Scientific Inc.). Slides were left to dry overnight in darkness, and then imaged on an inverted Nikon Ti microscope equipped with a CSU-22 spinning disk confocal. All images were captured with the following channel settings; DAPI at 300ms exposure time and 50% laser power, FITC at 300ms exposure time and 25% laser power, and CY5 at 100ms exposure time and 5% laser power. The images were overlapped using Fiji software (Schindelin *et al*., 2012).

### Radio-ligand uptake assay

The *in vitro* uptake assays were performed using methods developed in our laboratory as previously described (Yee *et al*., 2021; Koleske *et al*., 2022) (33759449, 36343260). After 48 hours of transient transfection of OCT1 variants in 96 well plates (poly-d-lysine coated), culture medium was removed, and cells were washed once with 250 µL HBSS. Four radiolabeled substrates of OCT1, [^3^H]-MPP^+^ (American Radiolabeled Chemicals, #ART 0150), [^14^C]-TEA (Perkin Elmer, #NEC298050UC), [^14^C]-metformin (Moravek, #MC 2043) and [^3^H]-sumatriptan (American Radiolabeled Chemicals, #ART 1619), were used in the assay. To each well, 80 µL of Hanks’ Balanced Salt solution (HBSS) (Gibco, #14025092) containing trace amounts of radioligand was added to each well and then incubated at 37 °C for 15 min. After a 15 min incubation period, the reaction mix was aspirated and the cells were washed 2× with ice-cold HBSS (250 µL). Next, 100 µL of MicroScint-20 (Perkin Elmer, #6013621) were added to each well and the plate was shaken at room temperature for 60-90 min. Radioactivity in each well was measured on a MicroBeta^2®^ Microplate Counter for Radiometric Detection (PerkinElmer). To determine the function of each variant, each variant was normalized to OCT1 reference and expressed as a percentage after background uptake measured in the GFP vector was subtracted from both, calculated as follows: (OCT1 variant minus GFP) *100/ (OCT1 reference minus GFP). Each variant was assayed in triplicate on a 96 well plate and measured in three or more biological replicates.

### Cytotoxicity assay

Cytotoxicity of SM73, a platinum analog, in OCT1 cells was adapted from our previous study (Zhang *et al*., 2006). To determine inhibition potencies of SM73 to kill 50% of the cells (IC_50_), cells were seeded on a 96-well plate (poly-d-lysine coated) at density 4000 cells/well. After 16-24 hours, the cells were incubated with different concentrations of SM73, starting from 100 µM to 1.7 nM, for 72 hours. Doxycycline (2 µg/mL) was incubated in the media with SM73 to induce the expression levels of OCT1. After 72 hours, media was removed and 50 µL of media was added to each well. Then 50 µL of Promega^®^ CellTiter-Glo^®^ luminescent reagent was added to each well. After 10-15 min incubation, transfer 80 µL to a 96 well plate (white, opaque). The luminescent cell viability was read on the Promega plate reader. The assay is based on quantification of the ATP present in each well, which relates to the number of viable cells in each well. The luminescent signal from cells treated with DMSO alone was considered the maximal signal (i.e. 100% cell growth). The percent cell growth of each OCT1 reference and mutants and at each concentration of SM73 were calculated. IC_50_ were determined using GraphPad (Prism v9.0).

For the deep mutational scanning, the cells (cells transfected with OCT1 library) were seeded into two T75 flasks. One flask only have doxycycline (2 µg/mL) and the other flask contained SM73 (1 µM) and doxycycline (2 µg/mL). After 48 hours, cells in both T75 flasks were split and transferred into new T75 flask and treated with doxycycline (2 µg/mL) with or without SM73 1 µM. Approximately 6 million cells were seeded in the T75 flask, and the T75 flask treated with SM73 1 µM, will have 10 million cells as may more cells will be killed after 48 hours. After another 48 hours, the above process was repeated. Every 48 hours, when the cells were split, approximately 1.5-3.0 million cells were collected for genomic DNA extraction. After another 48 hours, the above process was repeated. The cells in T75 flask are exposed to doxycycline (2 µg/mL) and SM73 1 µM or doxycycline (2 µg/mL) only for a total of 144 hours.

### Synthesis of SM73, a platinum anticancer agent

The methodology of Christen and Higgins was adopted for the synthesis of the 4-phenylpiperazinyl platinum analog SM73 (Giandomenico *et al*., 1995) (Supp Fig 16). The key species [PtIICl_3_NH_3_] was obtained as its potassium salt, K[Pt”Cl_3_NH_3_]^-^ (Supp Figure 13, Scheme 1 complex **1**) from cisplatin. Reaction of complex **1** with NaI followed by 1-phenylpiperazine afforded the mixed-halo Pt(II) complex **2**, which in turn was converted to the dichloride by formation of the aqua species with silver nitrate and the mixture exposed to HCl resulting in complex **3** (SM73). The synthesis of SM73 was previously described in a patent by Giacomini and More (Kathleen M. Giacomini, 2015). (SP-4-3)-Amminedichloro (cyclohexanamine) platinum (II) (Scheme 1, complex **3**, SM73). Synthesis of complex **1** was performed per the procedure described previously (Giandomenico *et al*., 1995). To an aqueous solution of **1** (0.375 g, 1.05 mmol) was added 0.4 g (2.0 mmol) of Nal in 1 mL of H_2_O and stirred for 30 min at ambient temperature. Then, 0.55 mL of 1-phenylpiperazine (0.1 mL, 0.58 mmol) was added to the reaction mixture and stirred for additional 4h at room temperature. The precipitated yellow solid was washed with water and ethanol. Additional washing with acetone provided compound **2** as a yellow solid, which was filtered and dried in vacuo (0.81 g, 52% yield).To a stirred suspension of **2** (0.50 g, 0.93 mmol) in 30 mL H_2_O was added 0.253 g (1.49 mmol, 1.6 equiv) of AgNO_3_ in the dark. The reaction was allowed to stir for 4h before being decolorized by activated charcoal. The precipitated AgCl was filtered and about 10 mL of concentrated HCl was added to the filtrate. The solution was left undisturbed at room temperature for 2 h and then at 4°C overnight. The product was collected by vacuum filtration, followed by repeated washings with water, ethanol and ether to obtain SM73 as a light-yellow solid (0.42 g, 54% yield). Anal. calcd for C_10_H_17_N_3_Cl_2_Pt: C, 26.98; H, 3.85; N, 9.44. Found: C, 26.82; H, 3.47; N, 9.41. ^1^H NMR (DMF-d_7_): 7.16 (2H, t), 6.90–6.73 (3H, m), 4.12 (3H, b), 3.58–3.31 (2H, m), 3.19–3.03 (4H, m), 2.84–2.62 (2H, m).

### OCT1 AlphaFold2 structure mapping

The AlphaFold2 predicted OCT1 structure was used for initial investigation (Jumper *et al*., 2021). We created scores for specific functional and abundance importance of each residue based on our DMS results. The folding importance score at a given position was defined as the sum of the absolute values of the GFP scores for variants at that position with GFP scores >= 0.68 (corresponding to loss of function) and is reported in Supplemental Table 14. The functional importance score for a given position was defined as the sum of the absolute values of the cytotoxicity scores for variants at that position with GFP scores < 0.68 (corresponding to non-loss of function) and is reported in Supplemental Table 15. These scores were mapped onto the B-factors of the AlphaFold2 model using the Bio3D R package for visualization (Grant, Skjaerven and Yao, 2021).

### Bioinformatics and sequence analysis

Conservation of OCT1 positions was determined using ConSurf (Yariv *et al*., 2023). Human and other specific sequences were obtained from UniProt (‘UniProt: the Universal Protein knowledgebase in 2023’, 2023). They were aligned with MAFFT using L-INS-i mode (Katoh and Standley, 2013). The alignment was visualized in Jalview (Waterhouse *et al*., 2009). For broader sequence analysis, first all sequences matching the SLC22 InterPro family (IPR004749) were downloaded (Paysan-Lafosse *et al*., 2023). Similar sequences in the unaligned set were removed by removing redundancy at 50% with Jalview, then the resultant set was aligned with MAFFT using L-INS-i mode. An HMM profile was generated from the alignment using HMMER3 (Eddy, 2011), then visualized with Skylign (Wheeler, Clements and Finn, 2014).

### Molecular Dynamics simulations

The Molecular Dynamics (MD) simulations of this work were done using GROMACS2022. The AlphaFold2 model of human OCT1 in an inward-facing open conformation was downloaded and prepared using the CHARMM-GUI membrane builder, where the ligands were parametrized using PDB coordinates and cGENFF. Partial charges were forced by QM cluster calculations in implicit solvent using the PCM model and were calculated using the 6-31G basis set with Hartree-Fock level of theory. Moreover, CHARMM-GUI was also used to prepare the mutated OCT1 simulations by automatically choosing reasonable rotamers, and for all OCT1 simulations to create disulfide bridges between three pairs of cysteine residues in the soluble extracellular domain. The systems consisted of OCT1 embedded in a POPC and solvated in a 10mM KCl solution with an ion imbalance such that the net charge was zero. The initial PBC box size was set to 90×90×100 Å^3^, and the systems were equilibrated according to the standard CHARMM-GUI protocol in the NVT ensemble. The production simulations were carried out using a 2 femtosecond timestep and a Nose-Hoover thermostat with separately coupled protein with ligand, POPC and water. For the barostat, the C-rescale scheme was used using a compressibility of 4.5e-5 and a tau of 5.0. The linear constraint solver (LINCS) algorithm was used to constrain all hydrogen-involving bonds, and particle mesh Ewald (PME) was utilized to calculate the long-range electrostatics above 12Å. For the LJ potential we used a neighbor-list cutoff with a switching function at 10Å and r_vdw_ of 12Å in consistency with the PME calculations.

The substrate MPP^+^ was docked using AutoDock with default parameters and a 10×10×10 Å^2^ box centered around the centre of mass of the entire transmembrane region. The top 100 poses were identical in most part thanks to the strong electrostatic forces.

### Enhanced Sampling MD Simulations

In this work, we used both non-equilibrium steered MD with umbrella potentials as an additional restraint in collective variable space, as well as the accelerated weight histogram method (AWH) to drive the conformational change. The expression for the CVs stemmed from the coevolutionary analysis (see below – Coevolution Driven Conformational Exploration) and was implemented in the GROMACS transformation pull coordinate module. For the calculation of free energy surfaces along two collective variables, we used the accelerated weight histogram (AWH) method as natively implemented in GROMACS. For all simulations, we used an adaptive target distribution with the value of 1 below the cut-off of 120kJ/mol and 0 above the cut-off with a smooth switching function between 100-140 kJ/mol. Additionally, we used the convolved bias potential to bias the reference coordinate and a harmonic coupling restraint between the reference and physical coordinates with varying force constants between 5.000-40.000kJ/mol, Å^2^.

AWH consists of two stages – the initial stage where the bias update size is decreased with each covering with the growth factor (set by default to 3.0), and after a criterion measuring the samples in the weight histogram and the actual number of collected samples is met the Wang-Landau algorithm is applied to achieve convergence. We recognized that our starting states may not be representative of the deepest minima, and thus we altered the growth factor to 2.0, which prolonged the initial stage such that more leeway would be given to account for time-dependent degrees of freedom to equilibrate. We could thus cover the conformational landscape several more times (<20) until the simulations exited the initial phase. We have heuristically seen that these estimations are more long-term reliable and produce flatter distributions without walkers getting stuck.

### Coevolution Driven Conformational Exploration

The core idea behind the computational methodology is to combine evolutionary information with physical information through machine learning. To achieve this, we further refined the procedure described in Mitrovic et.al 2022. To infer coevolving contacts between residue pairs, we chose the pre-constructed multiple sequence alignment of the PFAM transporter family PF00083. We then trained a Potts model using the direct coupling analysis (DCA) approach of the pseudo-maximum likelihood GREMLIN method, as described in Mitrovic et. al 2022. From the trained Potts model, we calculated the L2-norms of the pair-wise coupling parameter matrices. After average product correction (APC) and alignment to the target human OCT1 sequence, we could assign these coevolutionary coupling scores to all matched residue-residue pairs in the human OCT1 protein.

We then aligned the top 500 coupling pairs with the contact map extracted from the retrieved AlphaFold2 model of human OCT1, where we identified the positions presenting a high coevolution score but no inter-residue contact as so-called false positives (FPs). We then clustered the FPs according to coupling strength and created five separate coordinates for exploration with the accelerated weight histogram (AWH) enhanced sampling algorithm in GROMACS2022. After 5ns of sampling with 4 walkers sharing a bias, we trained a linear kernel SVM to distinguish the most extreme end states from the starting state. We could extract the top 50 distances and their coefficients and analogously generate a new CV set to be used in AWH again by retraining the SVM model with the new data. To avoid retaining previously identified and explored degrees of freedom we restrained the already sampled degrees of freedom through a 10-times higher regularization parameter. We repeated this process 5 times, each time increasing the sampling towards an outward open state. In total, we used 25ns per walker. These AWH simulations were run with a constant target distribution, and unrestrictive bounds. Using the last collective variable set, we defined bounds such that both start and end states would be sampled and set a cut-off of 120kJ/mol. We did not decrease the histogram size until the histogram equilibrated (meaning each point must have sampled within 80% of the target distribution at that point) and started two walkers from the start– and end states.

After achieving convergence of the free energy landscapes (Supp Fig 17), defined as the free energy surface that changes less than 1kJ/mol over 50 ns per point, we estimated the errors of the free energy surface by the procedure described in Mitrovic et.al 2022. In essence, we counted the transition imbalance between neighbouring points, which under equilibrium conditions and a well-estimated PMF, should be 0. By measuring the deviation from 0, we could estimate the size of the pointwise error in the free energy estimation (Supp Fig 18). Additionally, for a reliable free energy estimate, all walkers need to have sufficient overlap in their sampling regions. Thus, we measured the effective probability distributions after convergence and the overlap between walkers (Supp Fig 19)

### Mutational study

While 200-250 nanoseconds of simulation per walker were necessary to equilibrate, explore and converge the free energy landscape of the full conformational transition of the WT human OCT1, we instead focused our investigation of the mutants on the rocker switch mechanism, without sampling the opening of extra– and intracellular gates. We did this to alleviate the computational cost of running 18 mutants under 2 conditions each for a total of 36 separate simulation conditions. Under these conditions, we could iterate only 3 sets of explorations as described above. To enable a comparison between simulations, we used the optimal collective variable as derived for the WT system for the final AWH round converging the final free energy surfaces.

For each mutational condition, we visually inspected both the 3D structures from each basin to verify the functional assignment of each free energy basin, as well as the 2D landscapes themselves and quantitatively labeled each mutant based on this analysis on each.

### Protein network and computational importance analysis

While the collective variables inferred above may capture the essential degrees of freedom for between states, it is neither unique in its description, nor can it describe all of the 3N intricacie moving of the conformational change. Thus, after convergence, we prolonged the simulations (with a nearly static bias) until an additional 100 coverings of the conformational landscape were made. Based on this sampling, we obtained the frame-wise free energy estimate as a function of the collective variable:

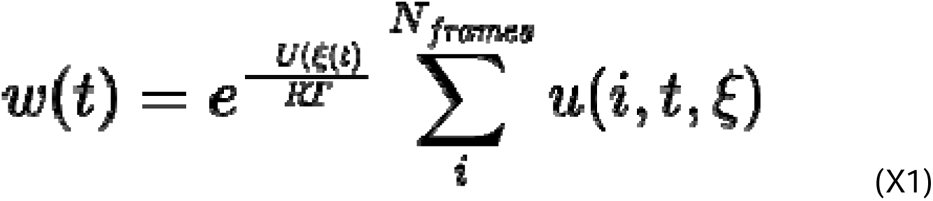

U is the pointwise free energy estimate, RT the thermodynamic constant at 298 K, and u(i,t,ξ) the binning procedure. Moreover, ξ(t) represents the value of the reaction coordinate at time t. Given a new reaction coordinate x, we could calculate each frame’s projection according to X2:

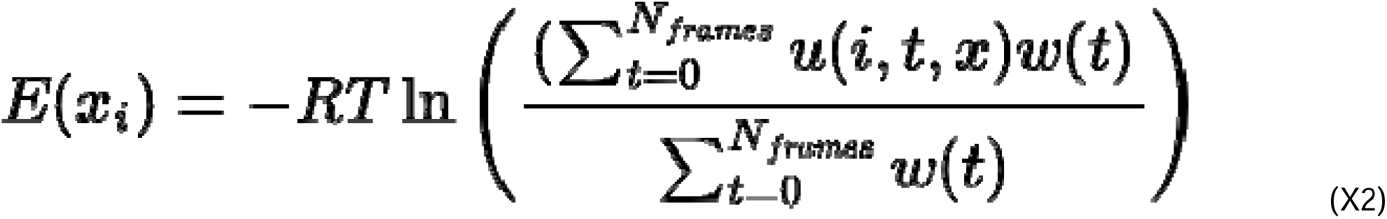

With this simple reweighting scheme accounting for the thermodynamic weight of each frame, we could reweight the free energy landscape onto any other degree of freedom. In this case, we systematically projected the free energy surface onto every minimum residue distance pair throughout the simulation. From these estimates, we measured the coupling between residues as the height of the highest free energy barrier along that coordinate, seeing as tightly coupled interactions should have two or more well-defined states with high barriers in-between, and less coupled pairs should have more spread-out distributions.

Additionally, we utilized the same reweighting procedure to back-map the 2D free energy landscape onto the determined SVM hyperplane normal vector, which would optimally describe the conformational change in 1D. This was primarily conducted for visualization purposes.

### Importance score profile peak assignment

With a robust conformation-wide measure of coupling, we constructed a protein network in which edges between residues are estimated as coupling strength mentioned above. We then analyzed the network and calculated betweenness centrality with the shortest-path finder algorithm as implemented in python’s networkx package. In simple terms, the quantity calculated is the number of shortest paths that pass through each node in the network. Given that the centrality values correlated well with the experimentally determined mutational sensitivity (as determined by a confusion matrix), we anticipated that we could possibly provide a molecular-level rationalization of the effect of mutations and we set out to identify which processes each of the peaks in the centrality plot were involved in. We focused on six different observable quantities, in particular residues which were primarily involved in:

1. Substrate dependency, determined by analysing the difference in centrality values between the apo and substrate bound simulations.
2. Extracellular communication, determined by measuring the percentage of shortest paths that connected the ECD and transmembrane domains.
3. The Rocker Switch mechanism, determined by mapping where the involved barriers localized on the free energy landscape. If the highest residue-residue pair barrier fell in the barrier region, we associated this with the main conformational change.
4. Gating contacts, determined by mapping the contacts to either the extracellular or intracellular gate opening and occlusion processes.
5. Inward-specific contacts, as residues involved in contacts only in the inward-facing states.
6. Outward-specific contacts, as residues involved in contacts only in the outward-facing states.

### Computational analysis of the biophysical basis of the effect mutations

Using visual analysis of the free energy surfaces themselves and 3D structures of each basin, we categorized each mutational condition into a functional phenotype that could be linked to the experimentally determined fitness (Figure 3). In detail, the categories we identified were:

1. High kinetic barrier, which was determined by measuring whether the kinetic barrier between inward– and outward facing states was more than 5kJ/mol higher than in WT simulations.
2. Low kinetic barrier, which was determined by measuring whether the kinetic barrier between inward– and outward facing states was more than 5kJ/mol lower than in WT simulations.
3. Destabilizes inward state, which was identified in cases where the inward facing state is rarely accessed compared to the outward facing state.
4. Disrupts substrate binding, which was identified in the case of E386K, where the substrate did not occupy the correct binding mode, rather it was the only condition in which the substrate would spontaneously unbind from the protein.
5. Stabilizes inward state, which was identified in cases where the inward facing state was more stabilized than the outward-facing state.
6. Substrate dependence loss, which was determined by examining whether new basins appeared, disappeared or changed in a significant capacity between substrate bound and apo simulations.

In some cases (I449T, S372G) no label could be attributed. The folding or trafficking-defect mutations were not probed computationally since a functional localization never could be obtained experimentally.

### GnomAD Database

To determine the frequencies of coding variants in human populations, we utilized the comprehensive gnomAD browser v2.1.1. Specifically, we extracted the variants for the SLC22A1 gene (transcript: ENST00000366963.4, NM_003057.3). We considered a variant to be functionally impaired if it had a score beyond two standard deviations of the synonymous variant distribution (Fig 1F). This corresponds to a cytotoxicity score of >= 0.68 or a GFP score of <= –0.72.

### UKBiobank total cholesterol, triglyceride and LDL levels

In this study, we utilize exome sequencing results available from ∼200,000 UK Biobank (UKB) participants (Bycroft *et al*., 2018; Szustakowski *et al*., 2021) to perform genetic association analysis of coding missense and in-frame deletion variants in SLC22A1 with total cholesterol and LDL levels of European ancestry and other ethnic populations. This research was conducted with approved access to UK Biobank data under application number 14105 (PI: J.S. Witte) and in accordance with the UK Biobank Ethics and Governance Framework. UK Biobank data are publicly available by request from https://www.ukbiobank.ac.uk. Individual exome sequencing data, LDL cholesterol and total cholesterol were extracted from the 199,933 UKB participants. The LDL cholesterol and total cholesterol levels were normalized using the R package bestNormalize (Peterson, 2021). The association analysis and rare variant analysis were performed using RStudio (2022.07.02). R package SKAT (version 2.2.4) was used for rare variant analysis (Wu *et al*., 2011). In SKAT analyses, variants that have a cytotoxicity score of 0.6798075 or higher, or an abundance score of – 0.7207321 or lower, are considered to have poor function.

We performed several association analyses to determine the significant association of OCT1 missense variant with total cholesterol and LDL levels and adjusted with body mass index (BMI), age, gender and for population structure (principal component analysis). The association analyses include:

1. Linear regression model to determine significant association of each missense variant with LDL and total cholesterol levels in European, South Asian and African populations (Suppl Table 6).
2. Combined the known poor function OCT1 missense variants with LDL and total cholesterol levels in European, South Asian and African populations. There are 5 variants that have poor OCT1 function in our study or from previous literature: p.R61C, p.C88R, p.G401S, p.M420del, p.G465R (Suppl Table 7).
3. Combined the common and less common poor function OCT1 missense variants that poor function and previously characterized: S29L, R61C, C88R, S189L, R206C, G220V, G401S, M420del, G465R (Suppl Table 7).
4. Performed SKAT analyses, including Burden test, which examines functional variants in SLC22A1 as causal factors with effects in the same direction (Suppl Table 8).

## Availability of data and materials

The sequencing analysis pipeline and data analysis code used in this publication is available in the github repository <repo link here>. Enrich2 scores are available as Additional file 2 and are also available at MAVE db <mave db link>. The sequencing data has been deposited at the NCBI Sequence Read Archive as BioProject PRJNA980726.

## Acknowledgements

We are grateful for you for taking the time to read our manuscript. We are also grateful to Matthew Howard, James Fraser, Aashish Manglik, Mary Hennessy, Anjali Jacob, Kliment Verba, Eric Greene, Margaux Pinney, Vijay Ramani, Robert Stroud, Nicholas Reyes, and for helpful feedback and discussion as we developed and conducted this project, as well as extensive guidance on the manuscript. We also would like to thank the hard work of those in the UCSF Flow Cytometry Core and Center for Advanced Technology without whom none of the FACS or sequencing could have been done.

## Ethics declarations

### Ethics approval and consent to participate

Not applicable.

## Author contributions

This study was designed by SWY, CM, DM, MK, LD, KMG, and WCM. SWY, MK, and WCM, did the preliminary experiments for deciding on conditions for the screen. SWY made the libraries and conducted the mutational scans. XZ performed the FACS sorting. CM and WCM processed the data from the mutational scan and mapped these onto the structures with support from DT. DB and SY with support from PRG made stable cell lines of mutants and conducted fluorescent microscopy. JY and SWY did the radioligand uptake experiments. SM synthesized SM73. SWY, CM, LK, and JSW did the population genetics analysis. DM and LD conducted modeling based on the AlphaFold2 states and conducted MD simulations. All authors contributed to writing the manuscript and making figures.

