## Supplemental Figures for "The full spectrum of OCT1 (SLC22A1) mutations bridges transporter biophysics to drug pharmacogenomics"

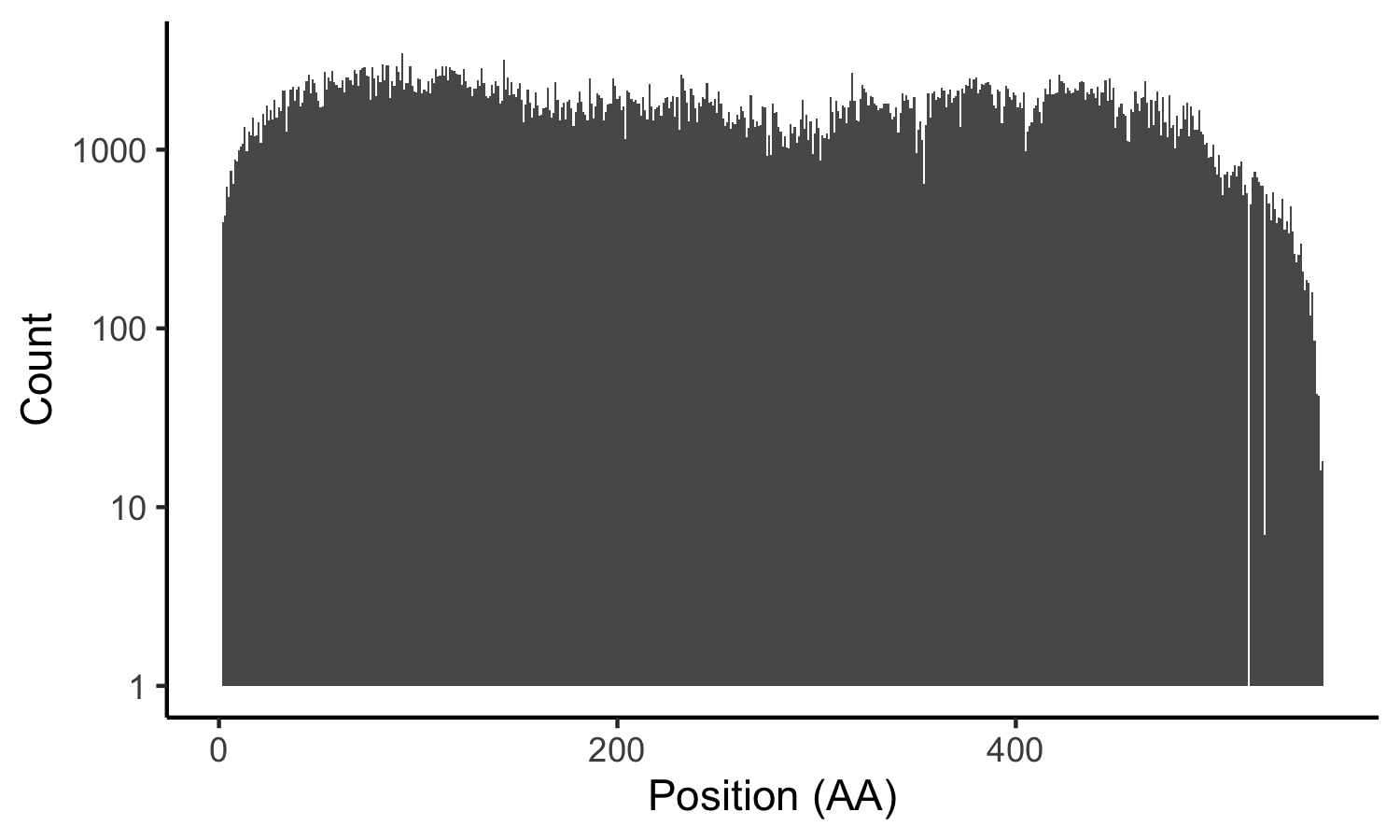

**Supplemental Figure 1. Baseline library counts in OCT1 gene.** Overall distribution of library counts (y axis) per positions (x axis) demonstrates the library is high quality with very few missing positions.

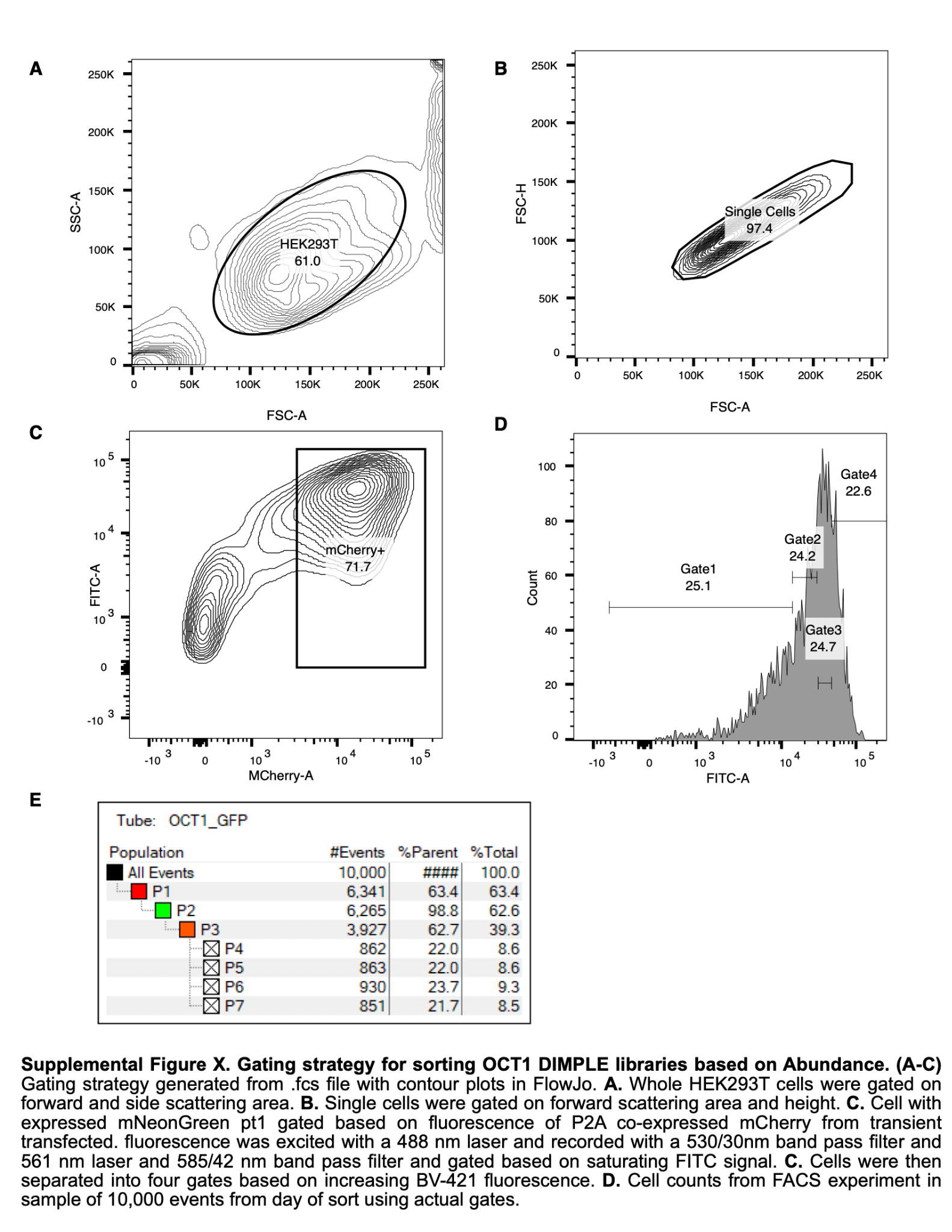

**Supplemental Figure 2. Gating strategy for sorting OCT1 DIMPLE libraries based on Abundance. (A-C)** Gating strategy generated from .fcs file with contour plots in FlowJo. **A.** Whole HEK293T cells were gated on forward and side scattering area. **B.** Single cells were gated on forward scattering area and height. **C.** Cell with expressed mNeonGreen pt1 gated based on fluorescence of P2A co-expressed mCherry from transient transfected. mNeongreen and mCherry fluorescence was excited with a 488 nm laser and recorded with a 530/30 nm band pass filter and 561 nm laser and 585/42 nm band pass filter and gated based once mNeonGreen fluorescence doesn’t depend on mCherry fluoresence. **C.** Cells were then separated into four gates based on increasing mNeonGreen fluorescence. **D.** Cell counts from FACS experiment in a sample of 10,000 events from day of sort using actual gates.

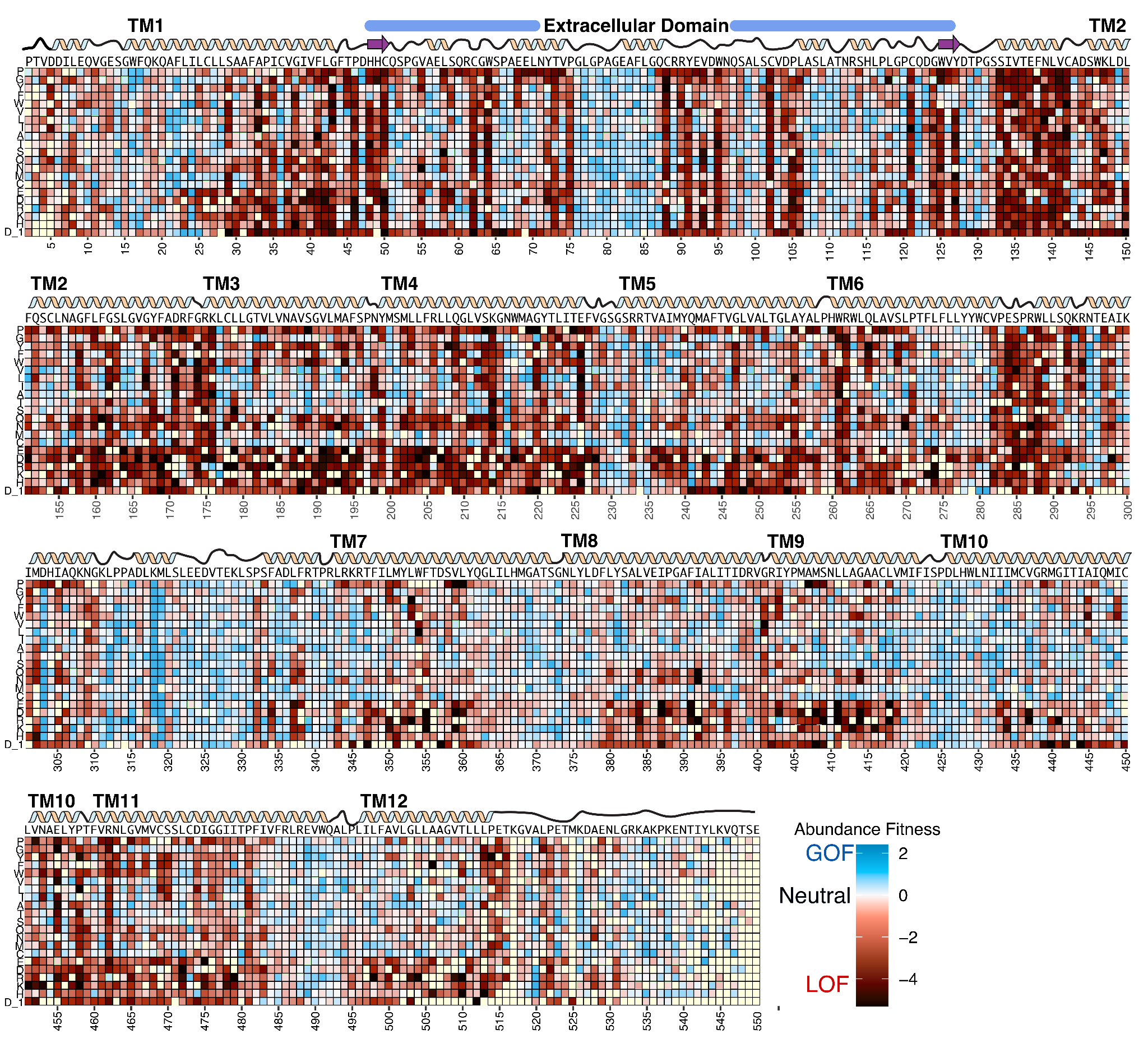

**Supplemental Figure 3. Abundance DMS Heatmap.** Abundance screen fitness effects depicted as a heatmap, with (x-axis) residue position versus (y-axis) mutation identity. In this assay, expression fitness colored from blue-to-red for increased to decreased expression relative to wildtype. Wildtype sequence of OCT1 is depicted above the heatmap with a cartoon of secondary structures as well. Missing data in light yellow.

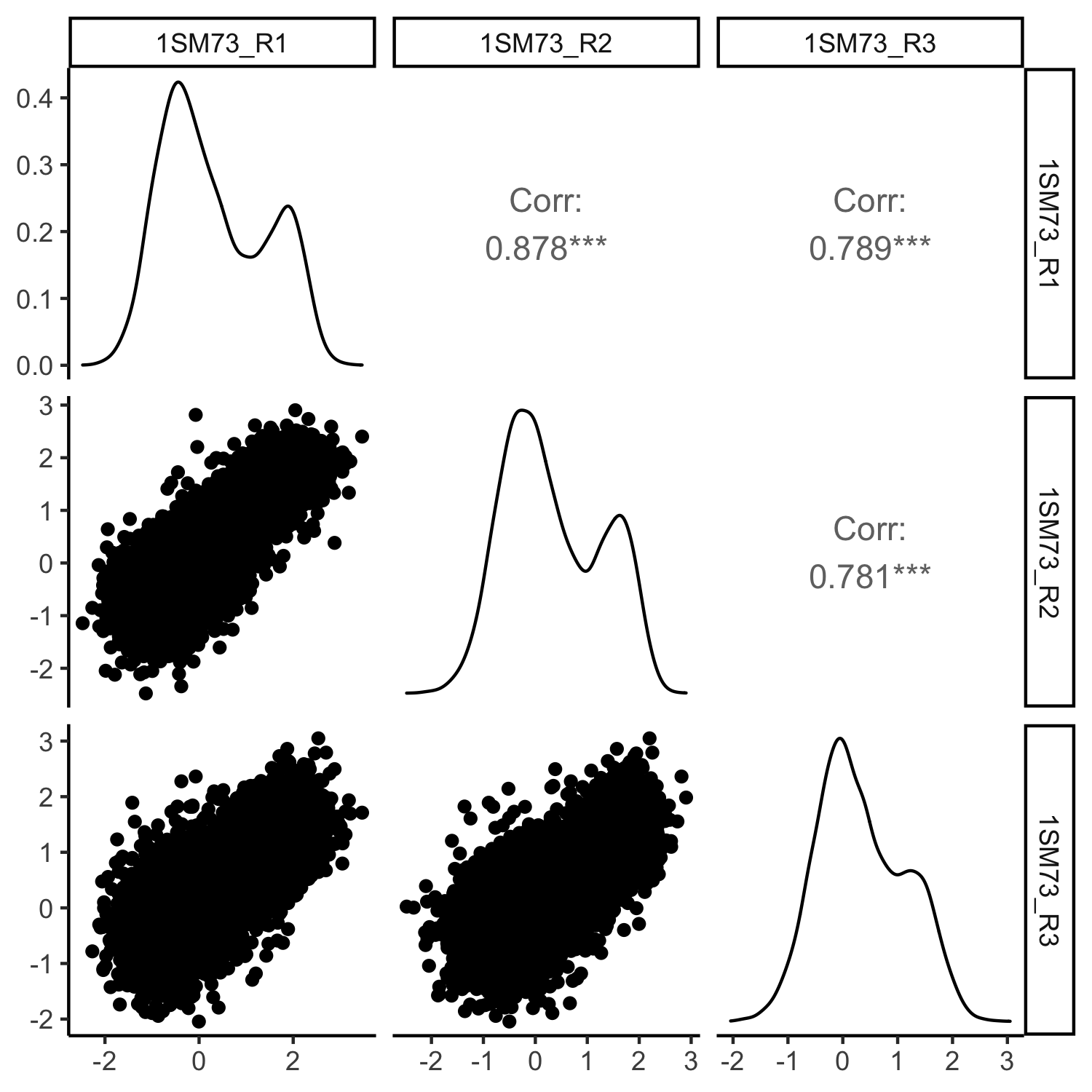

**Supplemental Figure 4. Correlations between three replicates of SM73 survival screen.** Cross-correlations between replicates, pearson correlation coefficients above the diagonal, with histograms of scores for replicates on the diagonal, and dot plots per varian below the diagonal. Overall, there is high replicability with replicate 3 being most different.

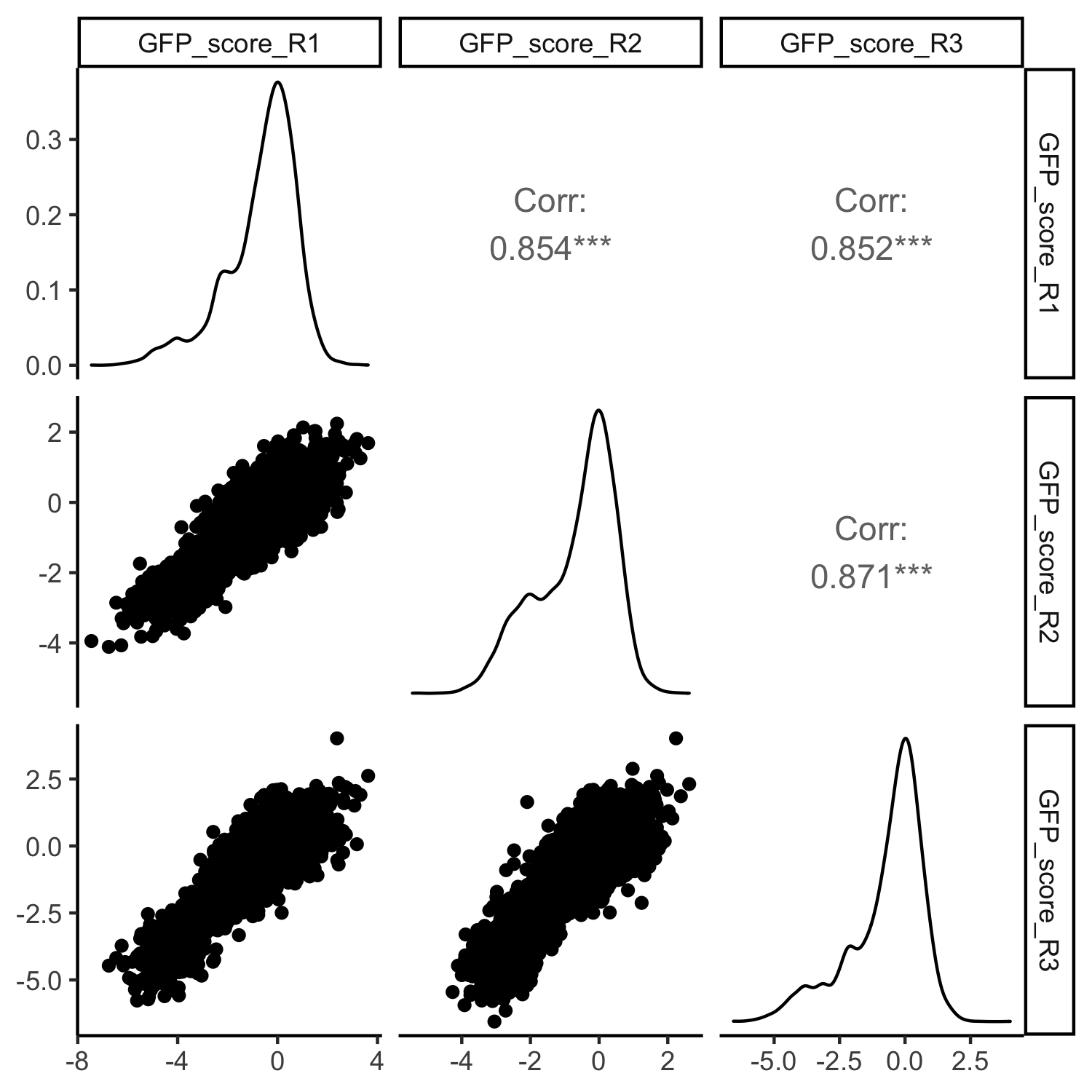

**Supplemental Figure 5. Correlations between three replicates of split fluorescent protein FACS expression screen.** Cross-correlations between replicates, pearson correlation coefficients above the diagonal, with histograms of scores for replicates on the diagonal, and dot plots per varian below the diagonal. Overall, there is high replicability with all replicates being highly similar.

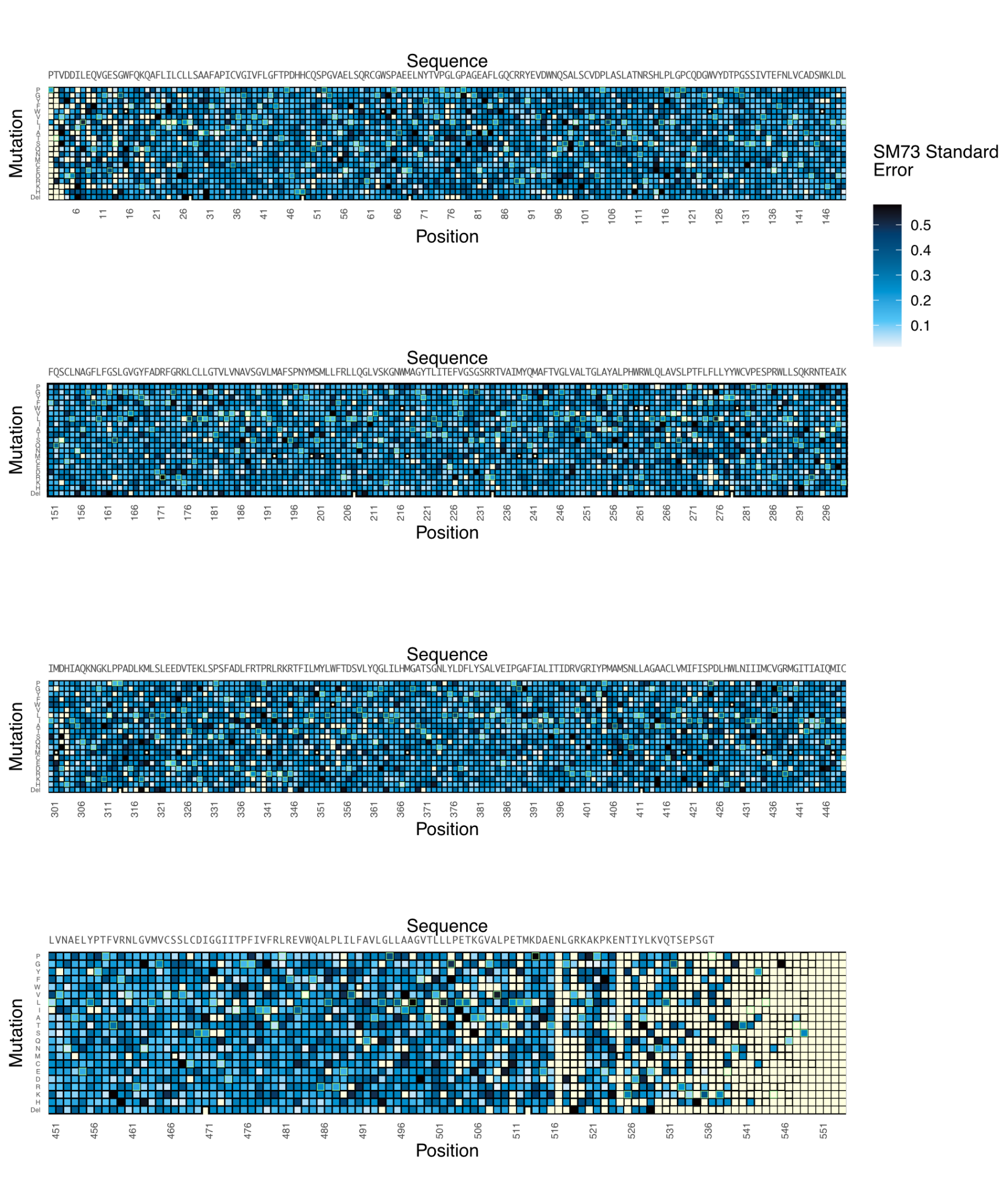

**Supplemental Figure 6. SM73 Standard Error Heatmaps.** Standard Error calculated per variant by Enrich2 in a heatmap with x axis position and y axis mutations.

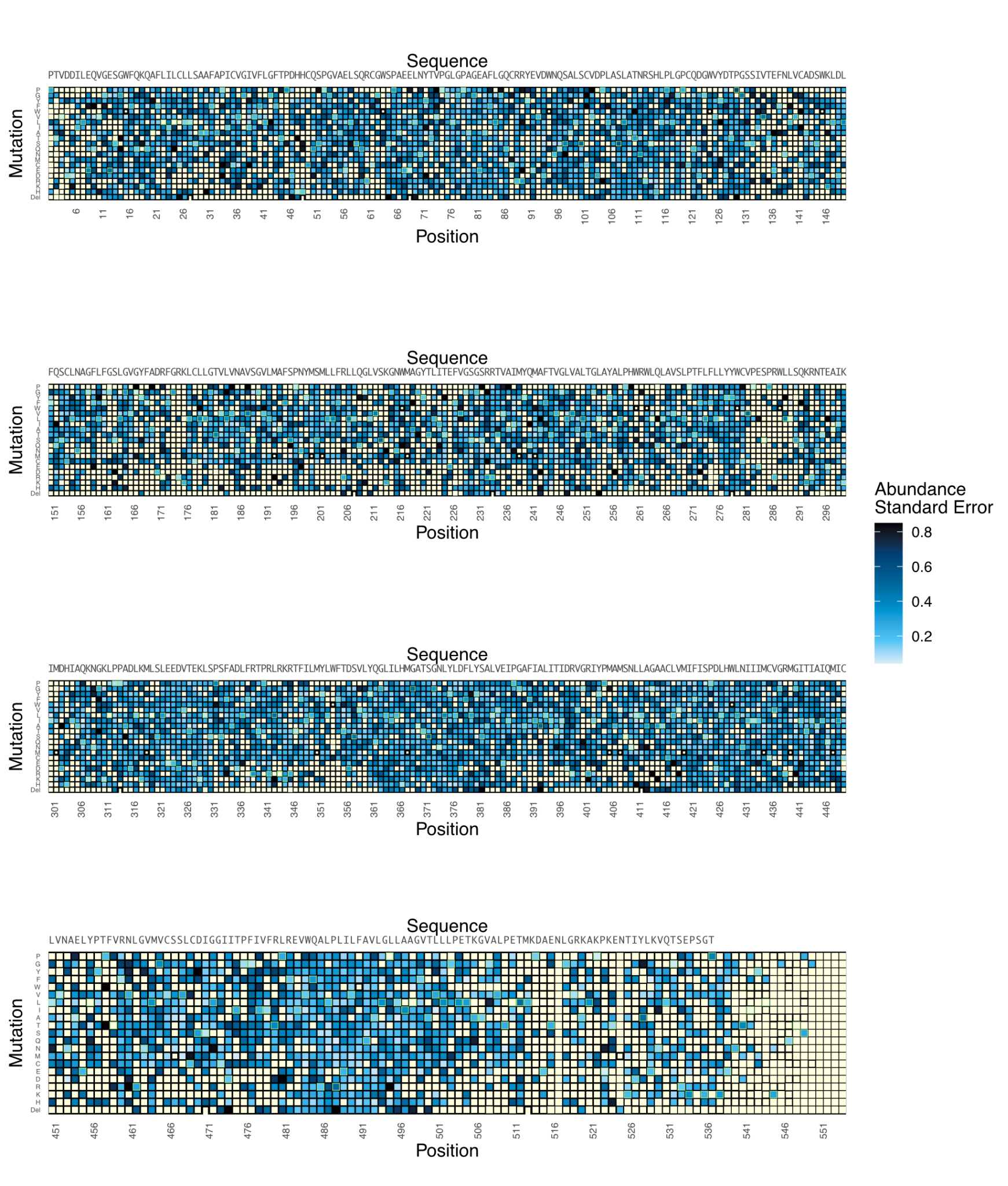

**Supplemental Figure 7. Abundance Fitness Standard Error Heatmaps.** Standard Error calculated per variant by Enrich2 in a heatmap with x axis position and y axis mutations.

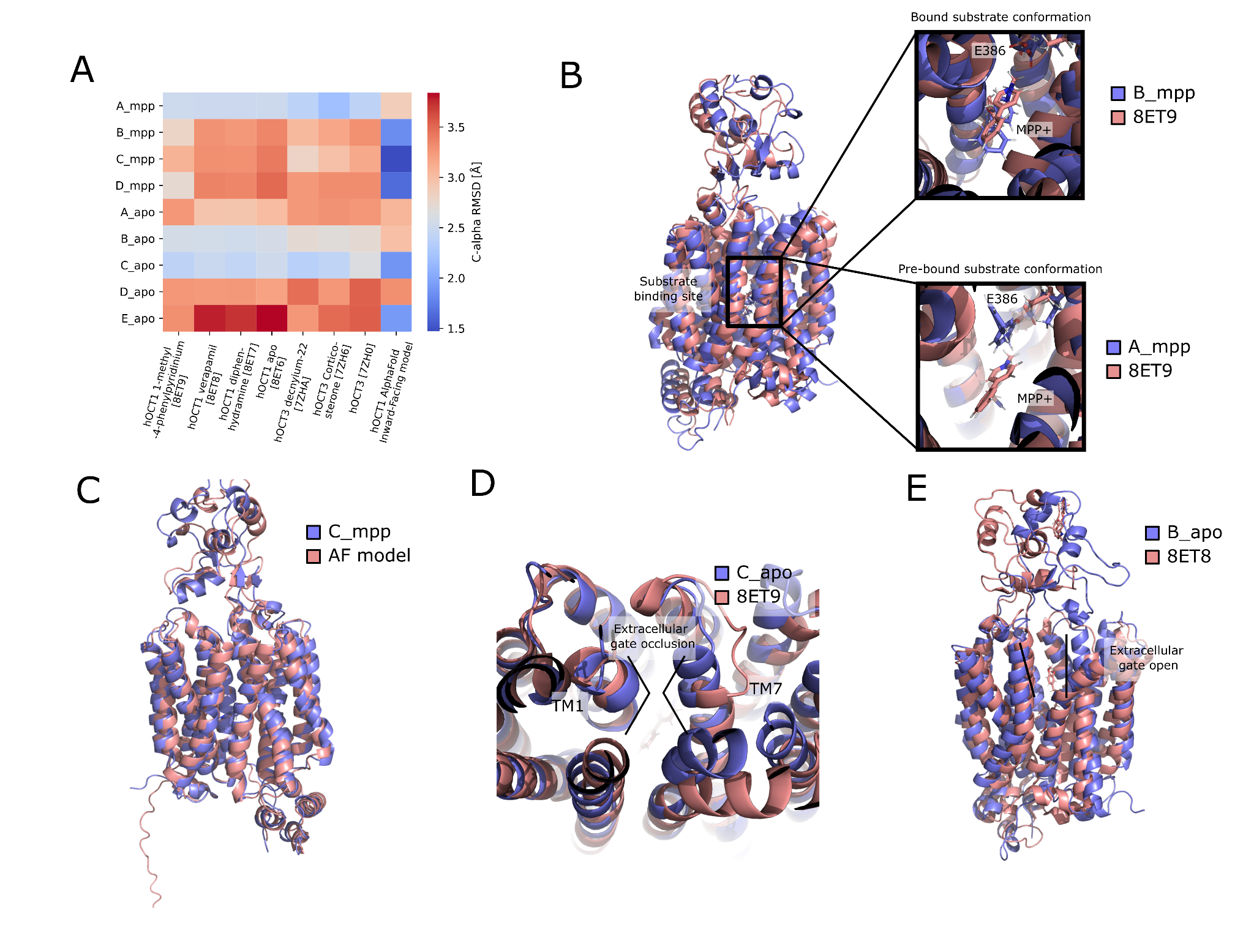

**Supplemental Figure 8: Structural Comparison between MD simulations and experimental structure determination**. A. Average C-alpha RMSD matrix between modelled basin snapshots (rows) and experimentally determined structures of OCT1 and OCT3 as well as the AlphaFold model generated in this work (columns). B. The substrate binding site and state-dependent coordination. In all basins but A_mpp the substrate is well coordinated and matches with the 8ET9 structure coordination. The A_mpp snapshots define a significantly more accessible substrate binding site.C. Good agreement between the energetically lowest substrate-bound basin and the Alphafold model. D. The structural details of the extracellular gate occlusion in the occluded basin match with the experimentally determined structure of OCT1. E. The open extracellular gate conformation of 8ET8 also matches well with the expectedly open B_apo basin.

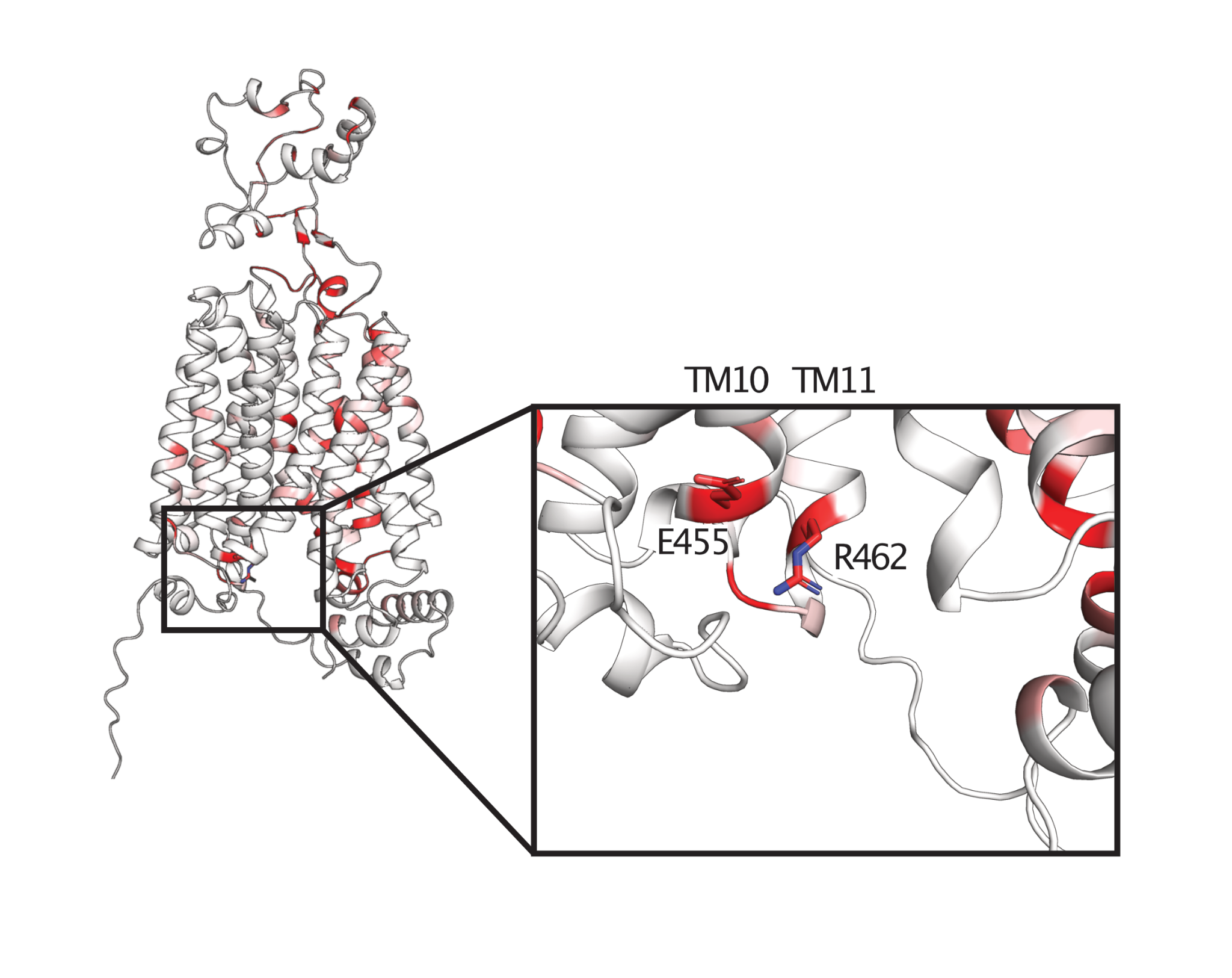

**Supplemental Figure 9: Interactions between the intracellular face of TM10 and TM11 drive stability.** Stability importance mapped onto the Alphafold2 based model in red based on the cutoffs shown in Fig 3A. Highlighted are residues E455 and R462 which likely interact during folding to stabilize folding. These residues are the main stabilizing residues outside of the C-terminal 6 transmembrane bundle that contribute to folding.

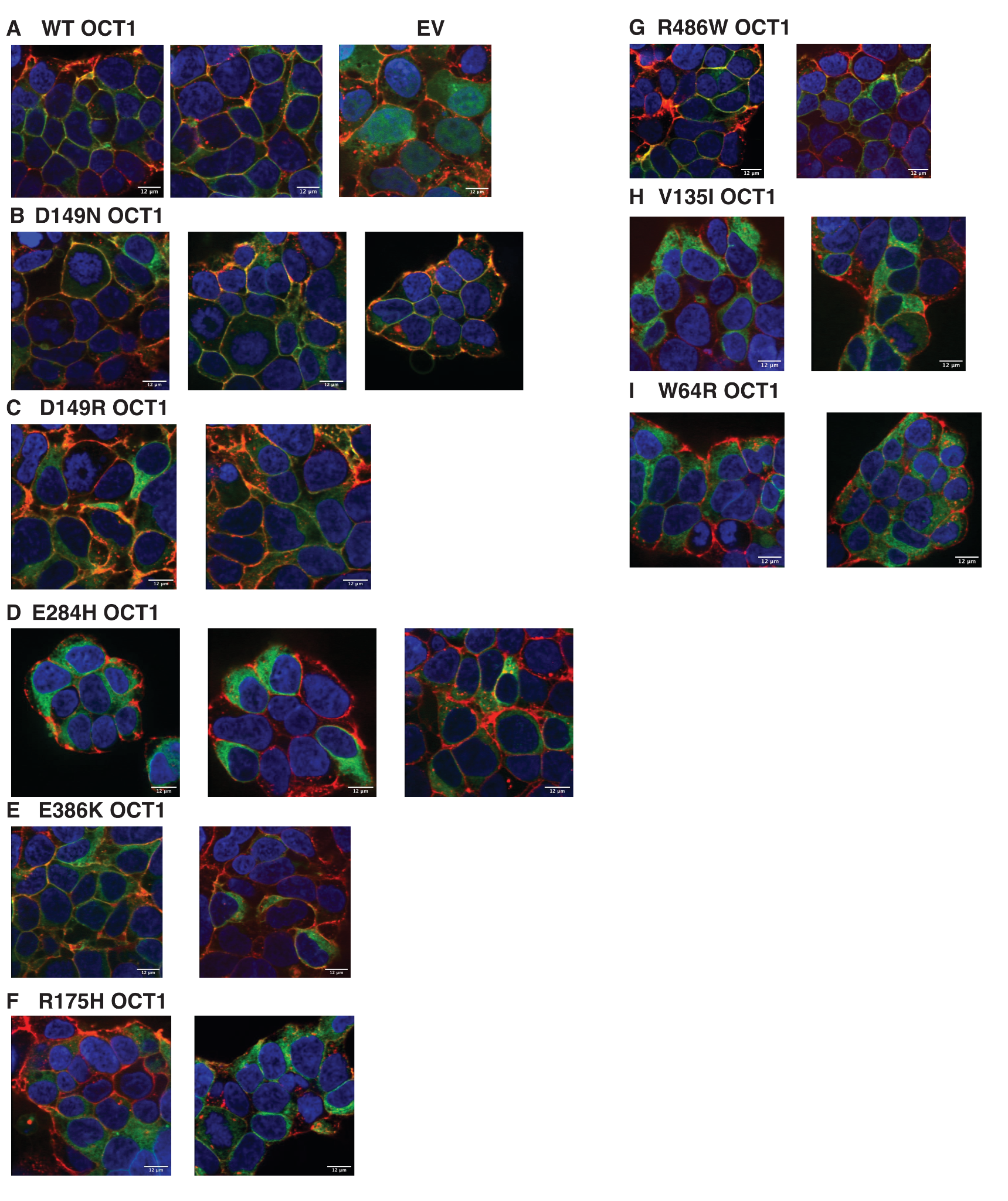

**Supplemental Figure 10: Confocal microscopy validation of OCT1 variant trafficking variants.** Confocal microscopy validation of trafficking phenotypes of **(A)** Wildtype and **(B-I)** variant expression patterns of variants which are loss of function based on abundance or cytotoxicity screen. Using EGFP-tagged OCT1 (green), cellular localization and expression are determined using nuclear stain (DAPI, blue) and cell surface stain (WGA-Alexa Fluor 647, red). Variants with low abundance identified in our screen (W64R, V135I, R175H, E284A) have commensurate low surface trafficking whereas WT and WT-like variants with high, loss of function, cytotoxicity scores (D149N, D149R, E386K, R486W) are properly expressed and trafficked to the surface. These results confirm trends seen within the high throughput screen.

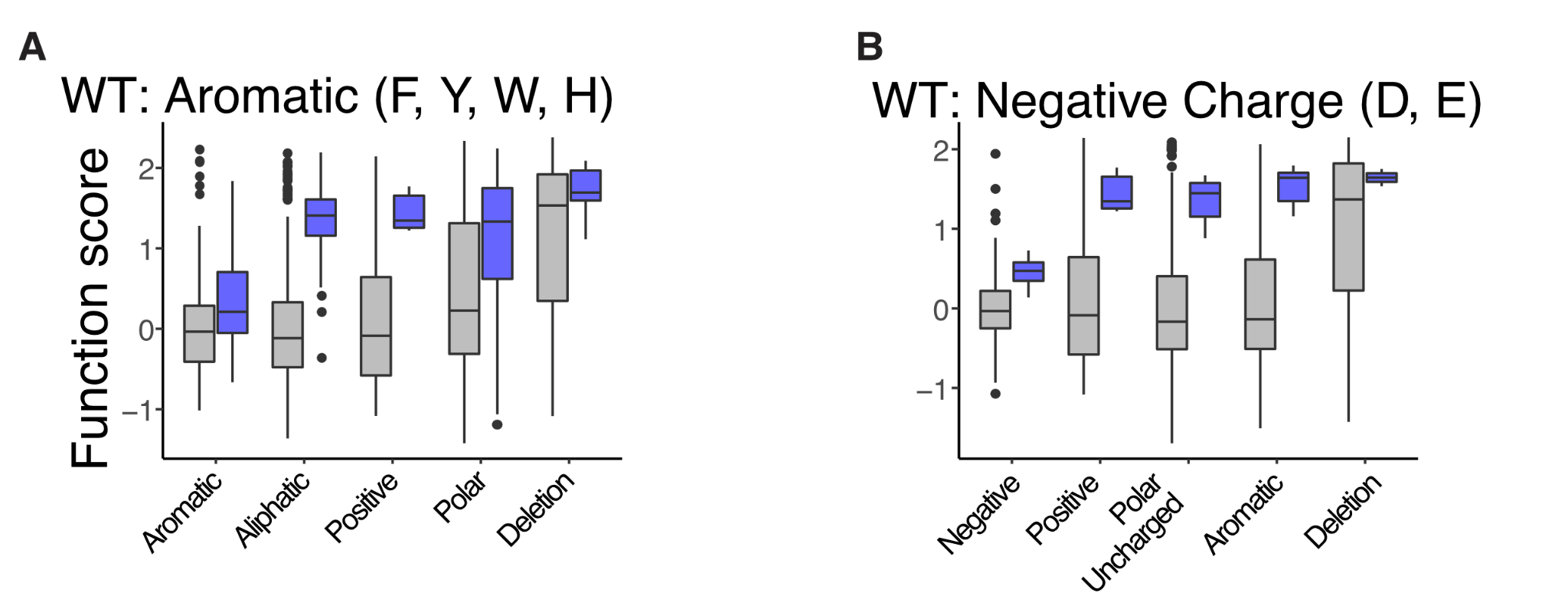

**Supplemental Figure 11: Sensitivity to physicochemical swaps within the substrate binding pocket.** Positions within substrate binding pocket (blue boxplots) show more sensitivity to changes of physical chemistry than other positions (gray boxplot): the functional scores of WT aromatic, or negatively charged residues show larger effects of changes to physical chemistry in the substrate binding pocket, characteristic of cation substrate binding. Substrate binding pocket residues are defined as being within 5 angstroms of the substrate in any of the MPP+ states.

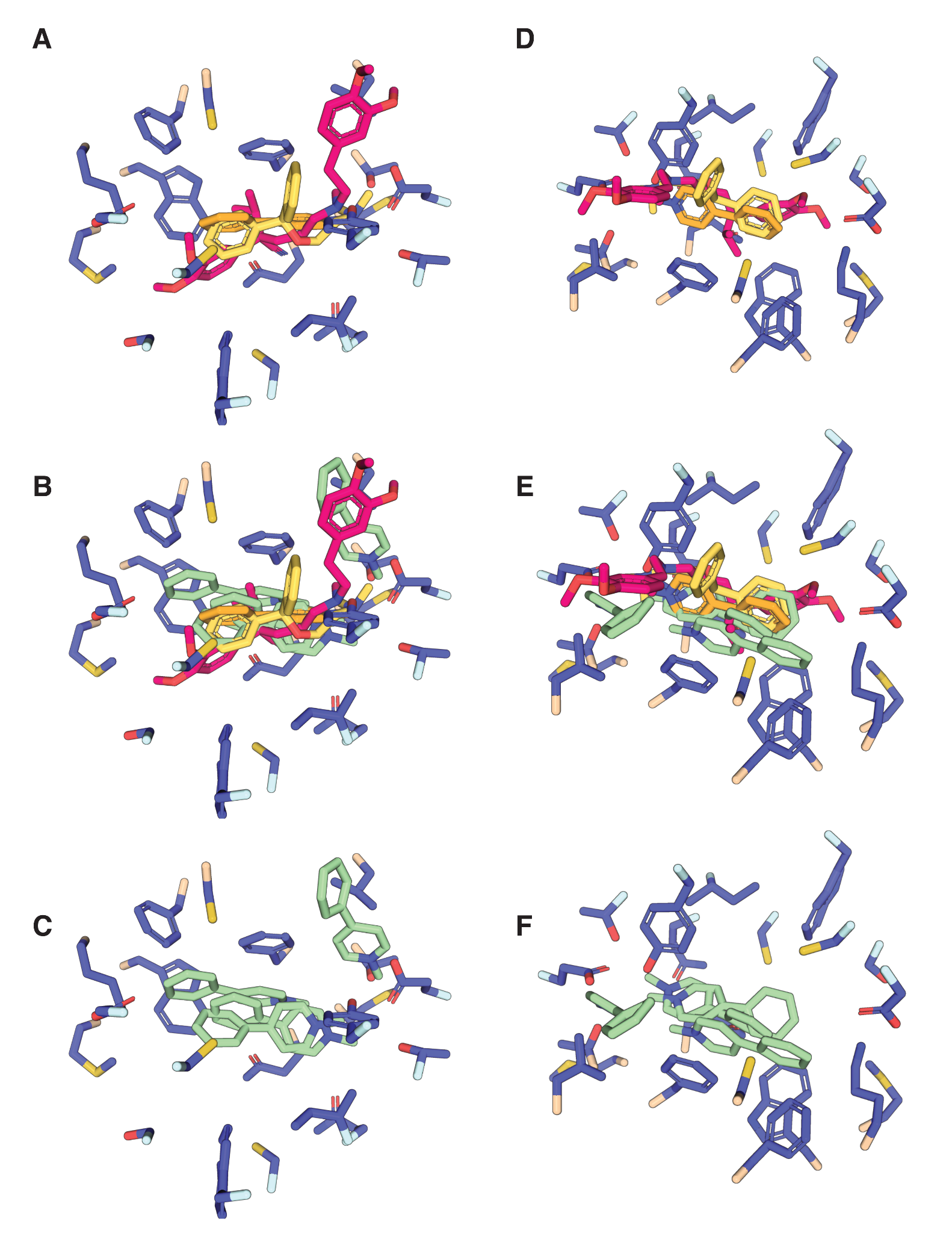

**Supplemental Figure 12: Ligand poses from experimental structures and simulations.** Comparison between ligand binding poses from predicted and structures from two perspectives **(A-C)** and (**D-F)** 180 degrees rotated from each other all with residues from MPP+-B predicted state. **(A, D)** shows experimental solved substrates diphenhydramine (yellow), pink (verapamil), and orange (MPP+) alone. **(B, E)** shows both experimental and predicted MPP+ positions (green). **(C, F)** Shows solely predicted states. Interestingly the one outlier state of MPP+ overlaps substantially with an aromatic from verapamil implying even that is a plausible location and likely represents an earlier binding site from MPP+ on the more outward facing face of the substrate binding pocket.

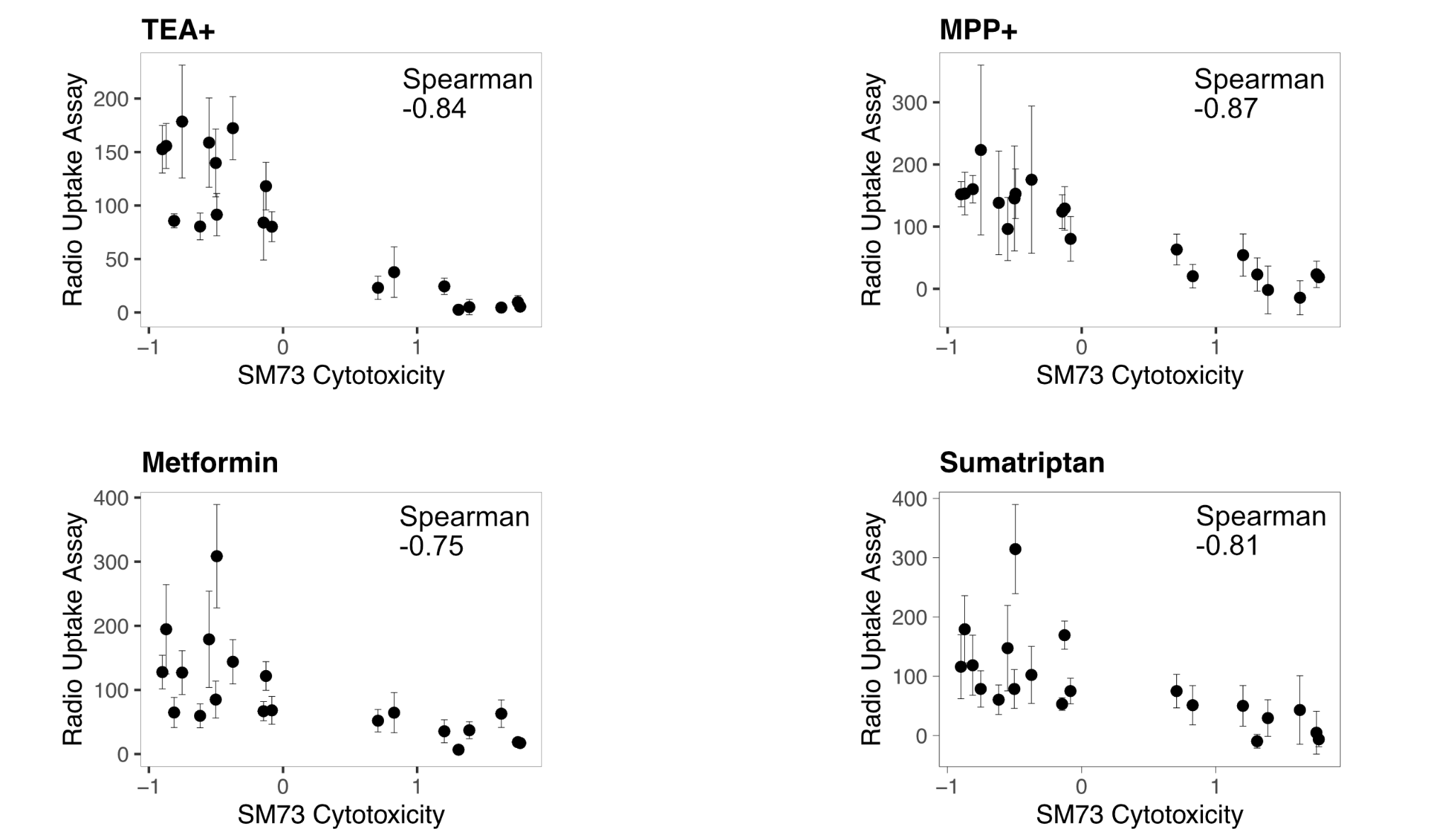

**Supplemental Figure 13: Correlation between cytotoxicity and validation radio ligand uptake assays.**Correlation between SM73 cytotoxicity fitness scores and radio-labeled substrates as uptake scores (error bars, SEM), with rank-order Spearman correlation coefficient shown.

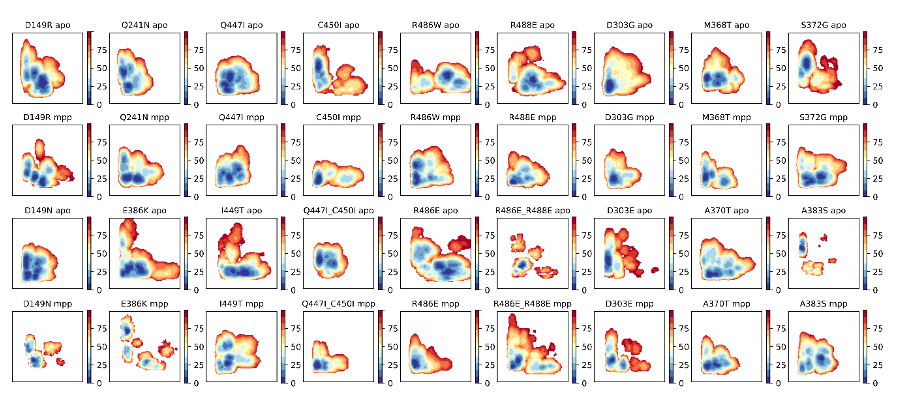

**Supplemental Figure 14: 2D landscapes of each mutation condition both with and without MPP+.** The colorbar signifies the energy level at each point in the surface, whereas each surface has a cutoff of 100kJ/mol to avoid showing energetically unfavorable regions. Note that due to micro-shifts in each conformation due to the mutation the collective variable space is not precisely transferable between different mutational conditions. We could however generally assign states based on the conformations found in each basin and the general position within the CV space. Within each condition the conformational states were separable and explorable.

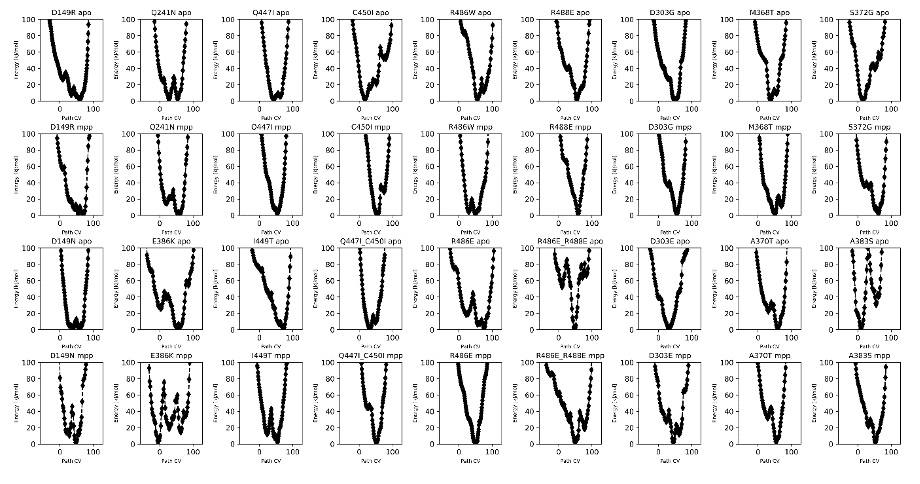

**Figure Figure 15: 1D landscapes of each mutation condition both with and without MPP+.** The landscapes were calculated by reweighting the 2D surfaces for each condition on the full collective variable taken as X-Y in the 2D landscapes.The Path CV signifies the progression along the resulting coordinate.

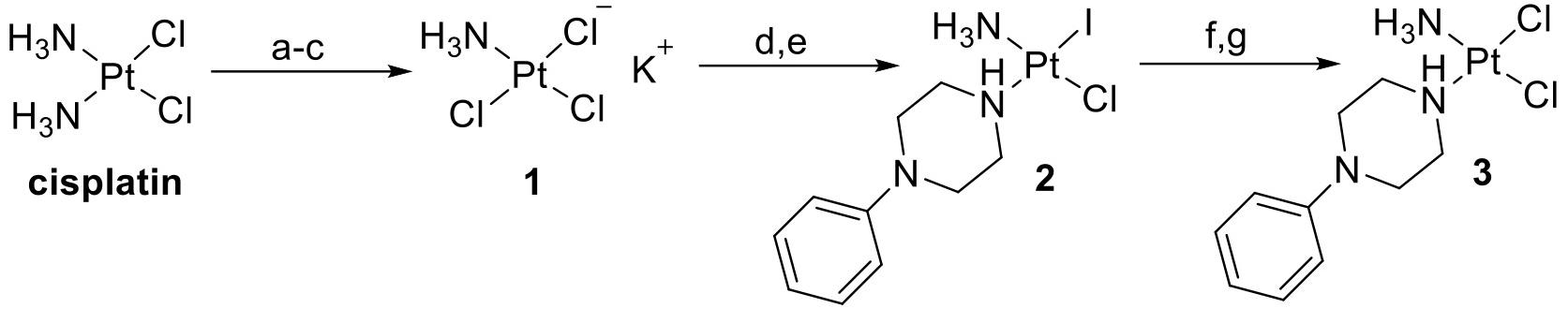

**Supplemental Figure 16. Scheme for SM73 synthesis.** Reagents and conditions. (a) Et_4_NCl, Δ, N_2_, dimethylacetamide, 8h, 100 °C; (b) Dowex 50W-X_8_ H^+^; (c) KCl, 4 °C; (d) NaI, H_2_O; (e) 1-phenylpiperazine, 4h, room temperature; (f) AgNO_3_, H_2_O, 4h, room temperature; (g) HCl.

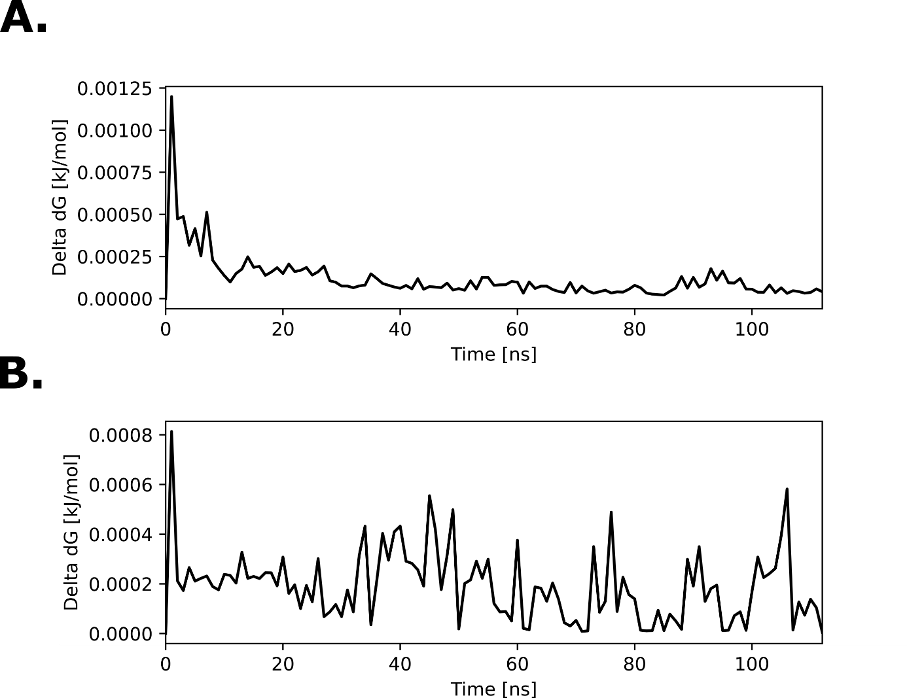

**Supplemental Figure 17: Molecular Dynamics simulation-wise convergence.** A. Behaviour of the Delta dG calculated every 10ns over time of the apo WT simulation. B. Behaviour of the Delta dG calculated every 10ns over time of the MPP+ WT simulation

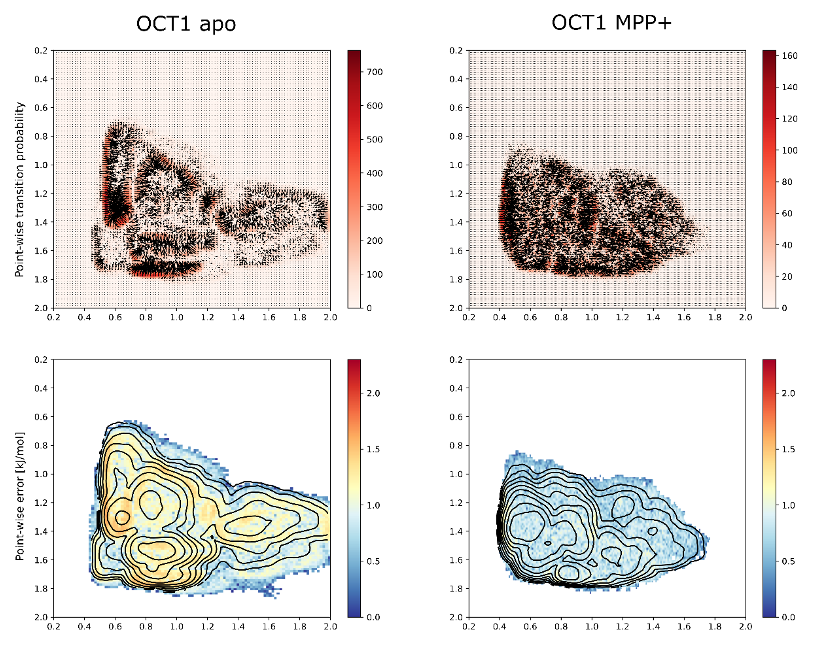

**Supplemental Figure 18: Simulation-wise transition probabilities and estimated error maps. Top.** The point-wise transition imbalances are represented as vectors and colored by the size of the corresponding transition matrix, where the units are raw counts. **Bottom.** To estimate the point-wise error we divided the transition between neighboring bins, from which we obtained a measure of the probability imbalance which in turn was converted to the free energy difference in kJ/mol. The error gives a local estimate of how far each point is estimated to be from a flat probability distribution. The black isocurves represent the free energy surfaces for the apo and MPP+ bound simulation conditions. See methods for details.

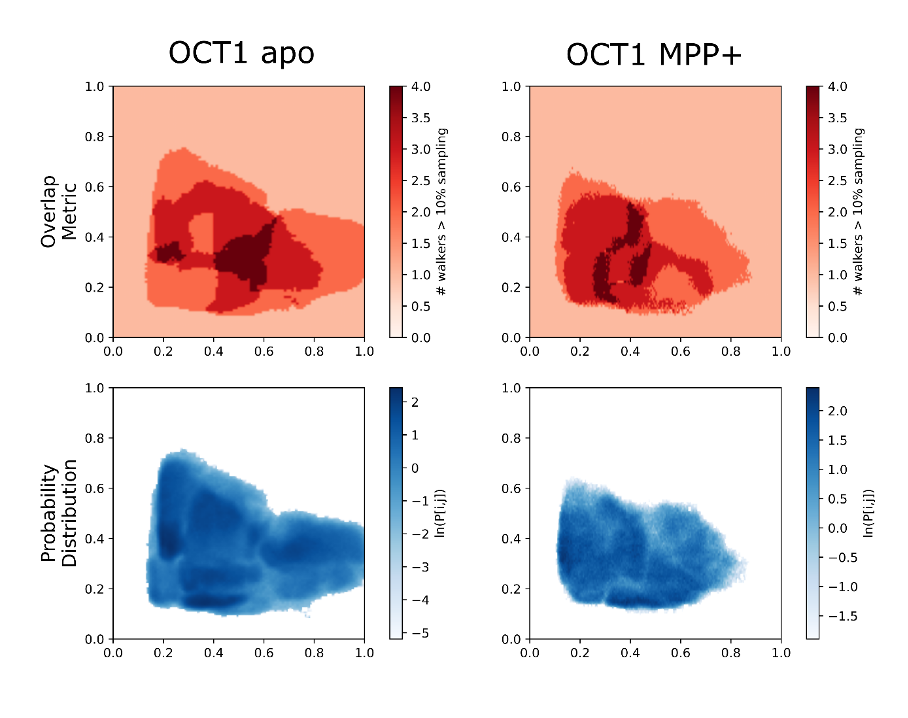

**Supplemental Figure 19. Simulation-wise probability and overlap maps for generating 2D landscapes**. **Top.** Biased probability distribution of each AWH simulation, aggregated from all walkers. **Bottom.** Overlap between walkers. Instead of calculating how many walkers ever visit each bin, we only summed walkers that had a probability of over 10% of visiting said bin. The color signifies this overlap metric.
